# A High-Affinity Nanobody Recognizing mNeonGreen Enables Versatile Biochemical, Cellular, and *in vivo* Applications

**DOI:** 10.64898/2026.06.30.735531

**Authors:** Maja Gere-Becker, Lisa-Marie Funk, Marie Gröger, Markus Kilisch, Pardis Najafi, Florian König, Federico Aloisi, Xuanfei Song, Lianne C. Davis, Antony Galione, Sabrina A. Hafeez, Benjamin L. Martin, Eugenio F. Fornasiero, Valeriia Kalienkova, Lisa Reinmuth, Petri Kursula, Hansjörg Götzke, Felipe Opazo, Steffen Frey

## Abstract

mNeonGreen (mNG) is among the brightest and most photostable monomeric green fluorescent proteins and is widely used for protein tagging. Here, we present sdAb(mNG), a high-affinity single-domain antibody (sdAb) that enables biochemical capture, imaging, and manipulation of mNG-tagged proteins. A 1.26 Å crystal structure reveals an extensive interaction surface between mNG and sdAb(mNG), accounting for its high affinity (K_D_ = 0.39 nM) and robust target recognition across diverse experimental conditions. This allows a single sdAb to support applications that typically require multiple specialized tools. We demonstrate the utility of sdAb(mNG) in several example applications including highly specific immunoprecipitation, direct immunofluorescence, and super-resolution imaging. Importantly, sdAb(mNG) retains high-performance target recognition even in intracellular environments. When expressed as an intrabody in living mammalian cells, sdAb(mNG) enables relocalization of mNG-tagged proteins to defined compartments or visualization of synaptic vesicle transport in primary neurons. In zebrafish, fusion of sdAb(mNG) to an F-box degradation domain induces cell-autonomous depletion of an endogenous mNG-tagged transcription factor and produces a clear developmental phenotype. These findings establish sdAb(mNG) as a versatile and robust affinity reagent that converts mNG from a passive fluorescent reporter into a multifunctional handle for imaging, proteomics, and programmable manipulation of endogenous and engineered proteins.

## Introduction

Genetically encoded fluorescent proteins (FP) have revolutionized cell biology by enabling the visualization and tracking of proteins in living cells. Among them, monomeric NeonGreen (mNG) has gained prominence for its exceptional properties, which outperform those of many established fluorescent proteins with similar spectral properties, such as GFP, EGFP, and mTagGFP. mNG exhibits significantly higher brightness (∼2.8-fold brighter than mEGFP) due to its superior quantum yield and extinction coefficient^1^. In addition, mNG matures rapidly (<10 min at 37°C versus ∼25 min for mEGFP), displays high photostability, and retains strict monomericity, properties that mitigate common limitations of fluorescent protein reporters, including aggregation, and aberrant protein localization in sensitive applications^2–4^. These characteristics make mNG especially well-suited for live-cell imaging, super-resolution microscopy, and quantitative fluorescence assays. To further expand the utility of mNG in biochemical and cellular workflows, target-specific high-affinity binders are required.

Single-domain antibodies (sdAbs) derived from camelid heavy-chain-only antibodies^5^ (also known as nanobodies) possess several advantageous properties, including small molecular size, high structural stability, and the ability to mediate strong, highly specific monovalent interactions. Their genetic accessibility facilitates the straightforward generation of fusion constructs with diverse functional modules, which can be expressed efficiently in prokaryotic and eukaryotic systems. Moreover, site-specific modifications, such as the introduction of ectopic cysteine residues, enable precise conjugation to fluorophores or solid supports. Collectively, these features render sdAbs powerful and versatile tools for affinity purification and advanced imaging techniques^6–10^.

Importantly, a subset of sdAbs remains functional in the reducing cytoplasmic environment, enabling their use in intracellular applications^11,12^. Genetically encoded sdAbs can, e.g., be fused to fluorescent proteins, enzymes, degradation tags, or localization motifs to create tools and sensors for target-specific imaging or manipulation in living cells or organisms. For example, fusion to degrons or E3 ligase adaptors can trigger ubiquitin-mediated degradation of tagged proteins^13,14^, while linkage to organelle-targeting sequences or scaffolding domains allows spatial redirection^15,16^ and synthetic rewiring of signaling pathways^17^. Such sdAb-based binders are thus ideally suited for use in inducible or conditional expression systems and offer precise, programmable control over the localization, abundance, and activity of their target proteins in living cells.

Here, we describe the identification and properties of a high-affinity sdAb specific to mNG. We report its binding characteristics, structural interface with mNG, and its application in several experimental workflows, including immunofluorescence, affinity-based protein pull-down, live-cell intracellular relocalization, protein tracking in primary hippocampal neuronal cultures, and *in vivo* target degradation in developing zebrafish. With this characterization palette, this work significantly enhances the available molecular toolbox, providing scientists with new options to visualize, track, tag, selectively degrade, or immunoprecipitate their favorite protein of interest fused to mNG.

## Results

### Generation and Selection of a Single-Domain Antibody Binding mNeonGreen

An alpaca immune sdAb-library was screened for binders recognizing recombinant mNG. Phage display panning yielded multiple positive clones, which were initially evaluated by single-clone ELISA. Candidate clones were further prioritized based on signal intensity and specificity in dilution ELISAs and on their performance in immunofluorescence assays using paraformaldehyde (PFA)-fixed cells expressing mNG fusions. The most promising clone 1E2 (Fig. 1A), designated sdAb anti-mNeonGreen (sdAb(mNG)), was expressed recombinantly in *E. coli* and purified for further biochemical and biophysical characterization.

**Figure 1:**
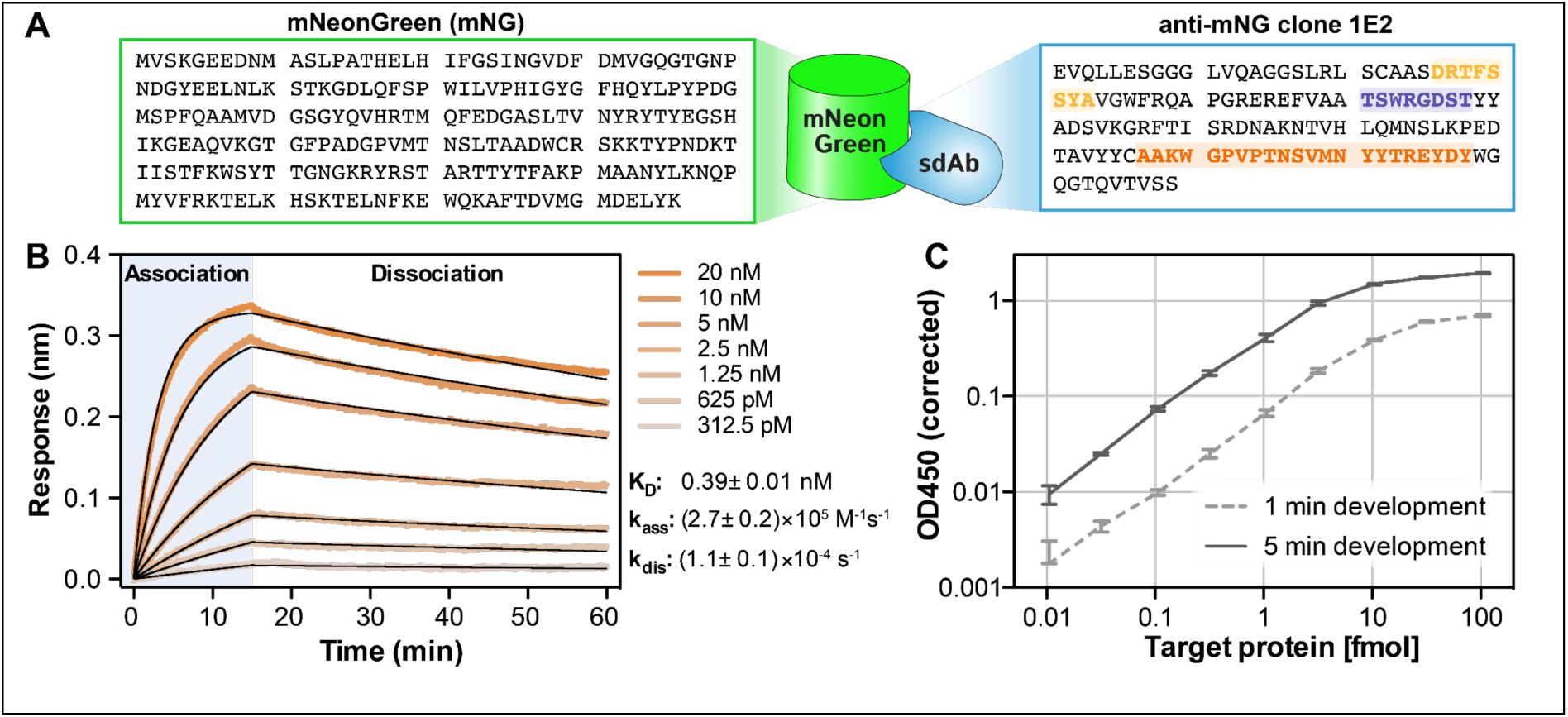
Characterization of the sdAb(mNG)/mNG interaction by BLI and ELISA. A: Sketch of the sdAb(mNG)/mNG complex (middle) with sequences of mNeonGreen (left) and sdAb(mNG) (right). CDRs 1, 2 and 3 of the sdAb (IMGT definition) are highlighted in light orange, purple or dark orange, respectively **B.** Binding kinetics of mNG to immobilized sdAb(mNG) measured by BLI. Soluble mNG was applied at varying concentrations, resulting in concentration-dependent association and slow dissociation kinetics. Global fitting of the data using a 1:1 binding model yielded an equilibrium dissociation constant (K_D_) of 0.39 nM, indicating high-affinity interaction. **C:** ELISA-based detection of immobilized recombinant mNG using biotinylated sdAb(mNG). The assay detected as little as 10 attomoles (∼0.3 pg) of target and showed a linear dynamic range over three orders of magnitude. Error bars represent standard deviations of three experiments performed in parallel.

### Characterization in BLI and ELISA

Binding affinity and kinetics were quantified using Biolayer Interferometry (BLI) (Fig. 1B). Binding of mNeonGreen to immobilized sdAb(mNG) yielded an association rate constant of 2.7*10^5^ M^-1^s^-1^, which is typical for sdAb-target interactions^6^, and a slow dissociation rate constant of 1.1*10^-4^ s^-1^. These values correspond to a calculated dissociation constant (K_D_) of 0.39 nM, indicating a highly stable interaction well suited for sensitive detection and affinity purification of mNG.

To assess its analytical performance in a direct ELISA format, sdAb(mNG) was site-specifically biotinylated via an ectopic cysteine and used to probe recombinant mNG directly coated onto ELISA plates. Detection with Streptavidin-HRP allowed detection of as little as 10 attomoles (∼0.3 pg) of recombinant target protein (Fig. 1C). Under non-limiting substrate conditions, the assay exhibited a linear dynamic range spanning approximately three orders of magnitude, underscoring the high sensitivity and quantitative capability of sdAb(mNG)-based detection.

### Structural Analysis of sdAb anti-mNeonGreen bound to mNeonGreen

The crystal structure of the sdAb(mNG)–mNeonGreen complex was solved at 1.26 Å resolution (Fig. 2, Supplementary Table 1). Consistent with the overall crystallographic model, synchrotron small-angle X-ray scattering (SAXS) measurements revealed a solution-state complex with matching size and overall shape (Supplementary Fig. 1 and Supplementary Movie 1). SdAb(mNG) recognizes a conformational epitope on the lateral surface of the mNeonGreen β-barrel, primarily involving β-strands 1, 2, 5, and 6 (Fig. 2A). The interaction engages all three CDR loops of the nanobody, forming an extensive, shape-complementary interface characterized by hydrogen bonding, several of which are mediated by water molecules (Fig. 2B, C), aromatic stacking (Fig. 2C, E), and van der Waals contacts. Of particular interest are several interactions involving β-strands 5 and 6 and primarily CDR3, as well as between CDR1 and CDR2 and β-strands 1 and 2. R111_mNG_ on β-strand 5 of mNeonGreen is engaged in a salt bridge with D117_sdAb_ and a π-π stacking with W100_sdAb_ of CDR3 in sdAb(mNG) (Fig. 2B, D). In the vicinity of this region, Y118_sdAb_ of CDR3 and Y32_sdAb_ of CDR1 form hydrogen bonds with Q124_mNG_ on β-strand 6 (Fig. 2B, D). Two pairs of aromatic residues (W53_sdAb_ on CDR2 and H18_mNG_ on β-strand 1, as well as Y32_sdAb_ on CDR1 and F20_mNG_ on β-strand 1) are engaged in aromatic stacking (Fig. 2E). Finally, K51_mNG_ on a loop succeeding β-strand 3 extends towards CDR1 and forms interactions with S30_sdAb_ and the backbone carbonyl of R27_sdAb_ (Fig. 2C). The sdAb’s N-terminus is in close proximity to mNeonGreen and participates in a hydrogen-bonding network (Fig. 2B), whereas its C-terminus extends outward, positioned approximately 4.5 nm from the mNeonGreen surface, an arrangement favorable for fusion or immobilization applications where steric freedom is beneficial. Similarly, the N- and C-termini of mNG are located away from the interaction surface, making them available for fusion-construct approaches.

**Figure 2.**
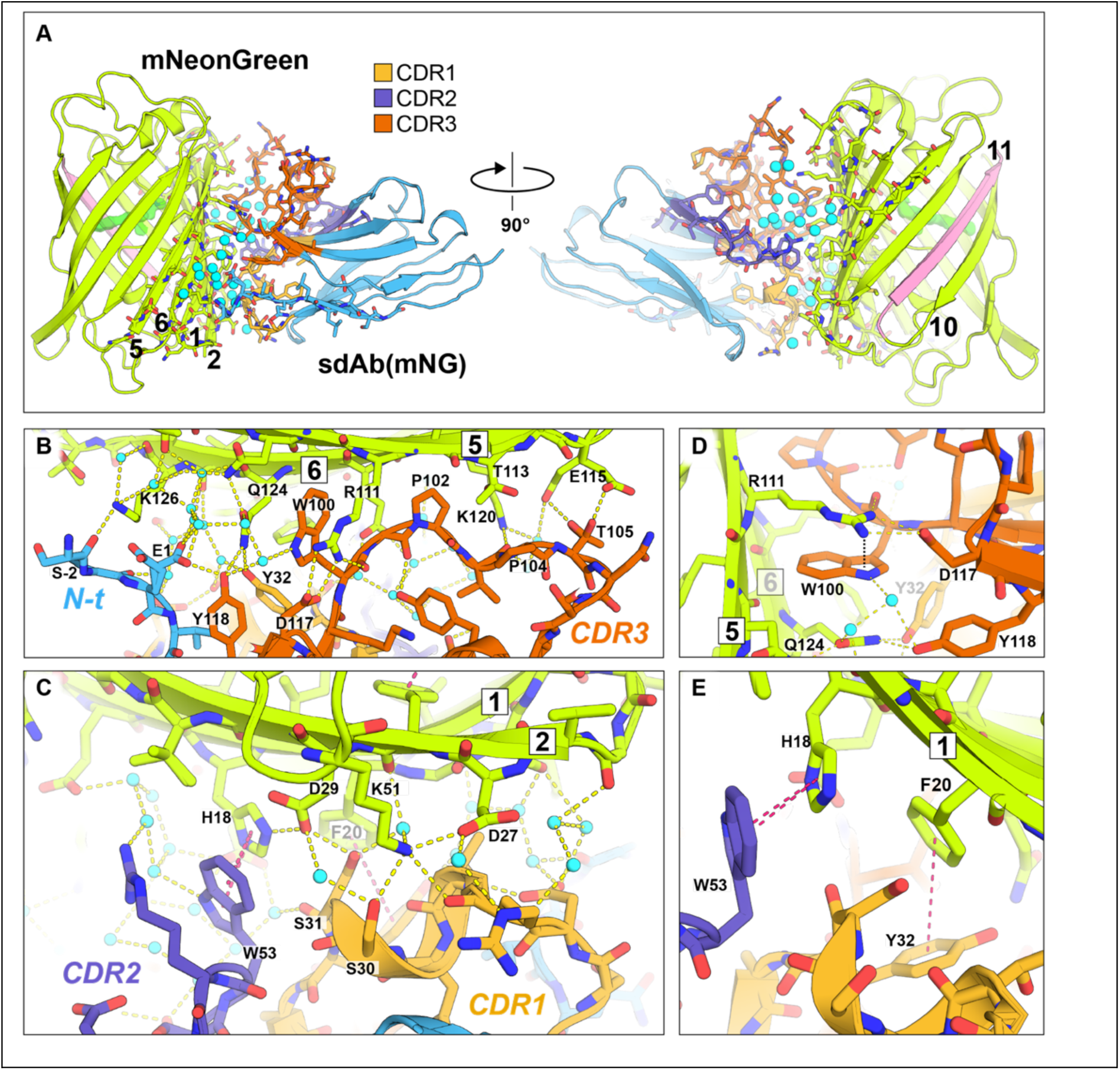
**Structural characterization of mNeonGreen and the sdAb(mNG) interactions (PDB: 30TF)**. **A**. mNeonGreen and sdAb(mNG) structure, CDRs of the sdAb are shown in unique colors. Water molecules filling the interface between the fluorescent protein and the sdAb and contributing to hydrogen bonds between the two proteins are shown as cyan spheres. mNG β-strand are numbered, interactions with sdAb(mNG) are mainly on β-strand 1,2 5 & 6. **B-E**. Close-up views of the interface between mNeonGreen and sdAb(mNG). Water molecules are displayed as cyan spheres, hydrogen bonds as yellow dashed lines, and π-π interactions as magenta dashed lines. Selected residues are labelled; mNeonGreen β-strands are indicated with boxed numbers.

### High-affinity capture of mNeonGreen-tagged proteins using immobilized sdAb(mNG)

To enable selective capture of mNG-tagged proteins, sdAb(mNG) was site-specifically immobilized on magnetic or non-magnetic agarose beads via a long flexible linker, generating high-affinity pulldown resins termed “mNeonGreen Selectors” (mNG Selector; Fig. 3A). Agarose-based resin (Agarose-magnetic too, but not shown) efficiently and specifically captured mNG fusion proteins from complex *E.coli* lysates with no detectable background (Fig. 3B, C).

**Figure 3:**
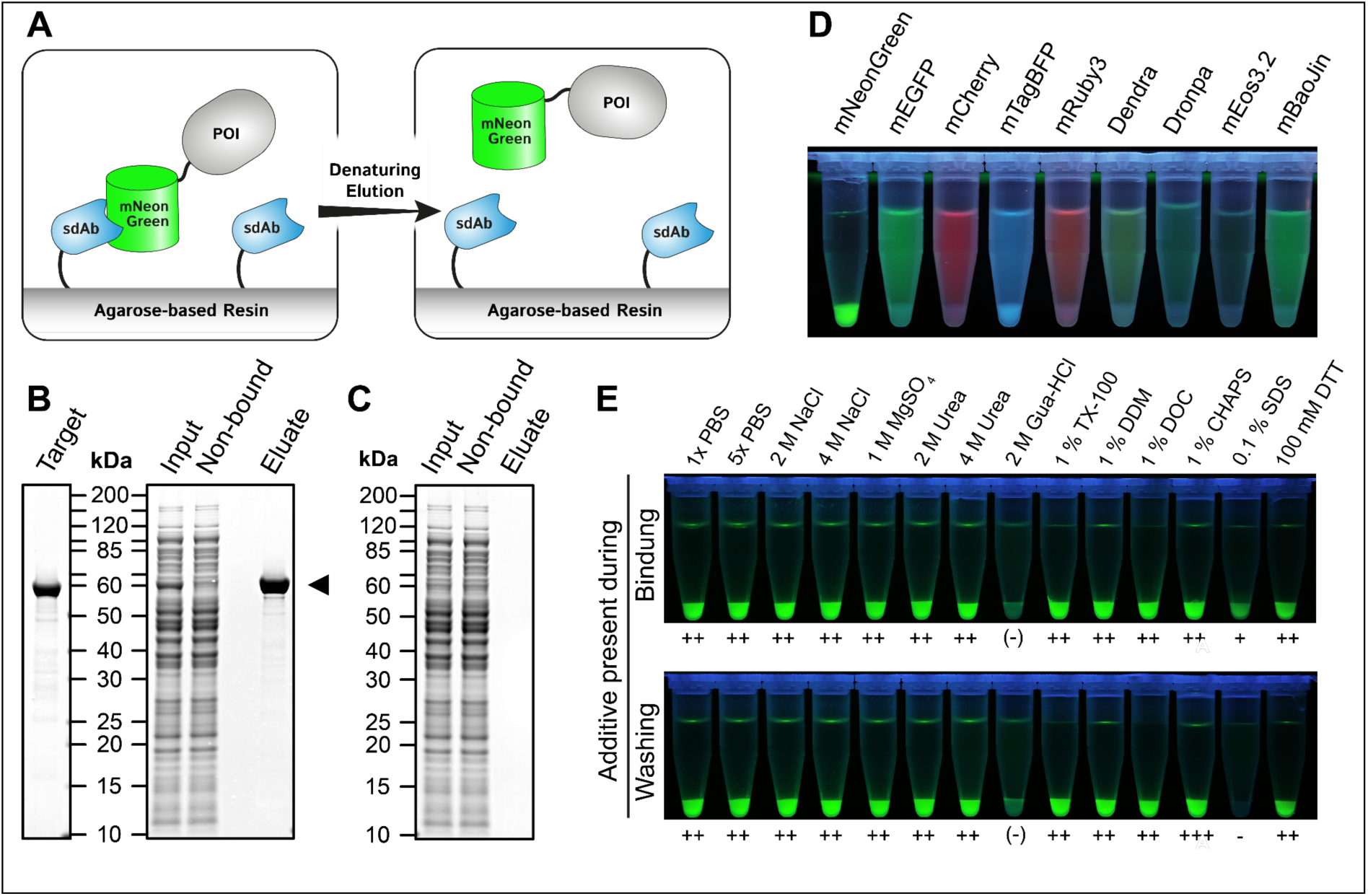
mNeonGreen Selector: a pulldown resin for mNeonGreen-fusion proteins. **(A)** Sketch of mNG Selector binding mNG fusion proteins. **(B, C)** Immunoprecipitation from *E.coli* lysate in the presence (B) or absence (C) of target protein (mNG-Halo fusion, arrowhead). Proteins were eluted in boiling SDS sample buffer. mNG Selector enables highly specific one-step capture of mNG-tagged proteins. (**D)** Binding assay to evaluate the cross-reactivity of the mNG Selector to other recombinant fluorescent proteins. Shown is a figure of the equilibrium bound fraction of each indicated fluorescent protein prior to buffer washing. (**E)** Compatibility of mNG Selector to various reagents applied during binding (top panel) or during washing steps (lower panel). mNG fluorescence is lost in the presence of 2 M Gua-HCl. The symbols below each row indicate binding efficiency: ++: strong binding, + binding,-: no binding.

To evaluate any potential cross-reactivity, equilibrium binding assays were performed using a diverse panel of fluorescent proteins from different evolutionary backgrounds (Fig. 3D). In this assay, mNG Selector demonstrated outstanding specificity, with no appreciable binding to any tested protein except for weak enrichment of mTagBFP under equilibrium conditions. This apparent binding to mTagBFP was completely lost after the resin was washed once with PBS (not shown). Notably, none of the residues of mNeonGreen engaged in direct contact with sdAb(mNG) are conserved in TagBFP (Supp. Fig. 2). To ensure this is a transient or irrelevant interaction, we again controlled the specificity of sdAb(mNG); this time, we performed an ELISA, confirming its selectivity for mNG, with negligible background signal from all other tested fluorescent proteins, including mTagBFP (Supp. Fig. 3).

Robustness of binding of mNG Selector to its target was assessed using a wide range of buffer additives (Fig. 3E). Capture was largely unaffected by high salt concentrations (up to 4 M NaCl or 1 M MgSO₄), non-denaturing detergents (1% Triton X-100, DDM, DOC or CHAPS), urea (≤4 M), or reducing conditions (100 mM DTT), regardless of whether additives were present during binding or washing steps. In contrast, binding was partially reduced or even completely abolished by low concentrations of SDS (≥0.1%) or by 2 M Guanidine-HCl present during the binding or washing steps, respectively. The interaction also remained largely stable down to pH 2.2, precluding conventional acid-based elution strategies (Supp. Fig. 4). These findings highlight the mNG Selector’s high tolerance for a broad range of biochemical conditions, with notable exceptions for strong denaturants.

**Figure 4.**
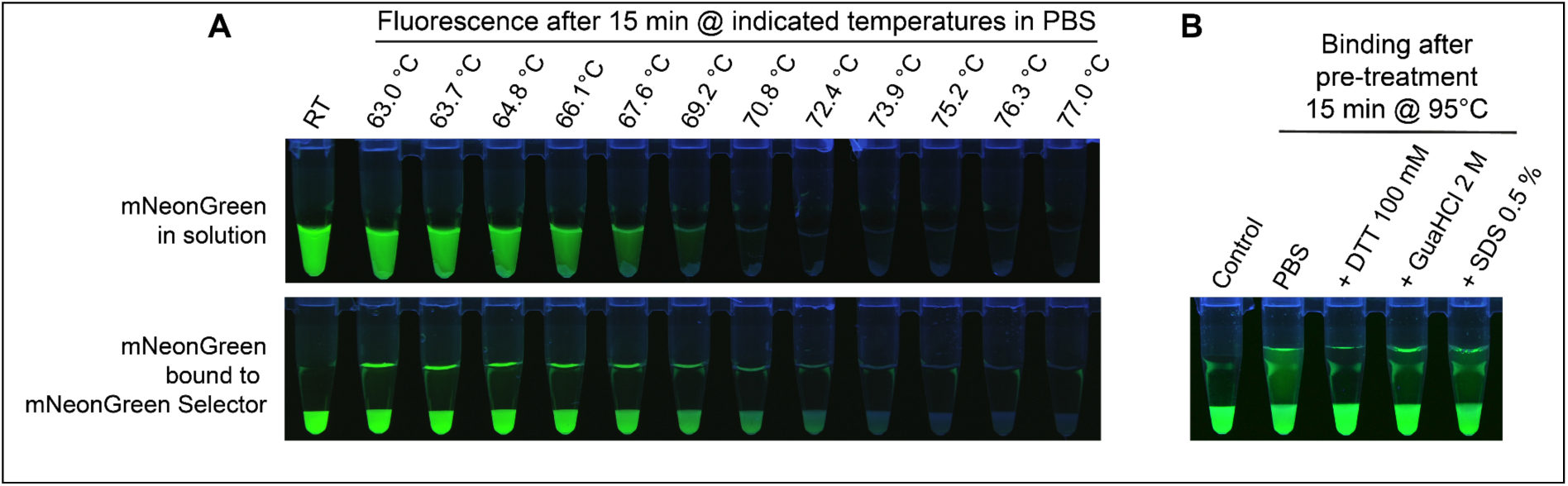
Temperature stability of mNeonGreen. (**A**) Thermal stability of mNeonGreen alone (upper panel) and in complex with immobilized sdAb(mNG) (mNeonGreen Selector; lower panel). Free mNeonGreen shows loss of fluorescence and precipitation upon incubation at increasing temperatures, whereas mNeonGreen bound to mNG Selector remains stable at higher temperatures, indicating stabilization by sdAb(mNG). (**B**) Thermal resilience of immobilized sdAb(mNG). mNG Selector was subjected to heat treatment under various conditions, including reducing and denaturing environments, followed by cooling and assessment of binding activity. Substantial binding capacity was retained under all tested conditions, demonstrating high thermostability and refolding capability of sdAb(mNG).

We next compared the thermal tolerance of the sdAb(mNG)-mNG complex with that of mNG alone. Free mNG exhibited visible loss of fluorescence and partial precipitation already after 15 min at 63°C, with complete loss at 70.8°C (Fig. 4A). When bound to mNG Selector, unfolding was shifted to higher temperatures, with fluorescence loss beginning only above 66°C and becoming nearly complete above 73.9°C, indicating that the binding of the sdAb(mNG) stabilizes its target.

As no temperature-induced release of fluorescent mNG from the mNG selector was observed, the data further suggest that sdAb(mNG) is more thermotolerant than its target. To directly assess sdAb thermostability, mNG Selector was incubated for 10 min at temperatures up to 85°C. After cooling down to room temperature, all samples retained full binding activity (not shown). In a more stringent test, mNG Selector was heated for 15 min to 95°C in PBS alone or PBS containing 100 mM DTT, 2 M guanidinium hydrochloride, or 0.5% SDS (Fig. 4B). After cooling and washing, substantial binding capacity remained under all conditions, with the highest recovery observed after heating in the presence of DTT. These experiments demonstrate that immobilized sdAb(mNG) can spontaneously refold into a functional state after near-complete thermal denaturation, even in the absence of an intact intradomain disulfide bond.

### Immunofluorescence detection of mNeonGreen-tagged proteins

For direct immunofluorescence (IF) applications, sdAb(mNG) was engineered to contain two ectopic cysteines for site-specific chemical conjugation of fluorophores (X2). The resulting conjugates (FluoTag-X2 anti-mNG) were validated in IF assays using PFA-fixed mammalian cells expressing mNG, localized to the surface of mitochondria (TOM70), or nuclear pore complexes (NUP98) (Fig. 5A & Supp. Fig. 5A, respectively). In these assays, FluoTag-X2 anti-mNG conjugated to Alexa Fluor 647 (AF647) was shown to robustly detect mNG without nonspecific background (Fig. 5A). To enable conventional indirect IF workflows using secondary reagents, recombinant sdAb(mNG) fusions harboring either the rabbit IgG or the mouse IgG1 heavy chain Fc antibody domain were generated (rHcAb(mNG)_Rabbit_ or rHcAb(mNG)_Mouse_, respectively). These Fc-fusion reagents were used as primary reagents, followed by species-specific secondary sdAbs for detection (Fig. 5B & Supp. Fig. 5B).

**Figure 5.**
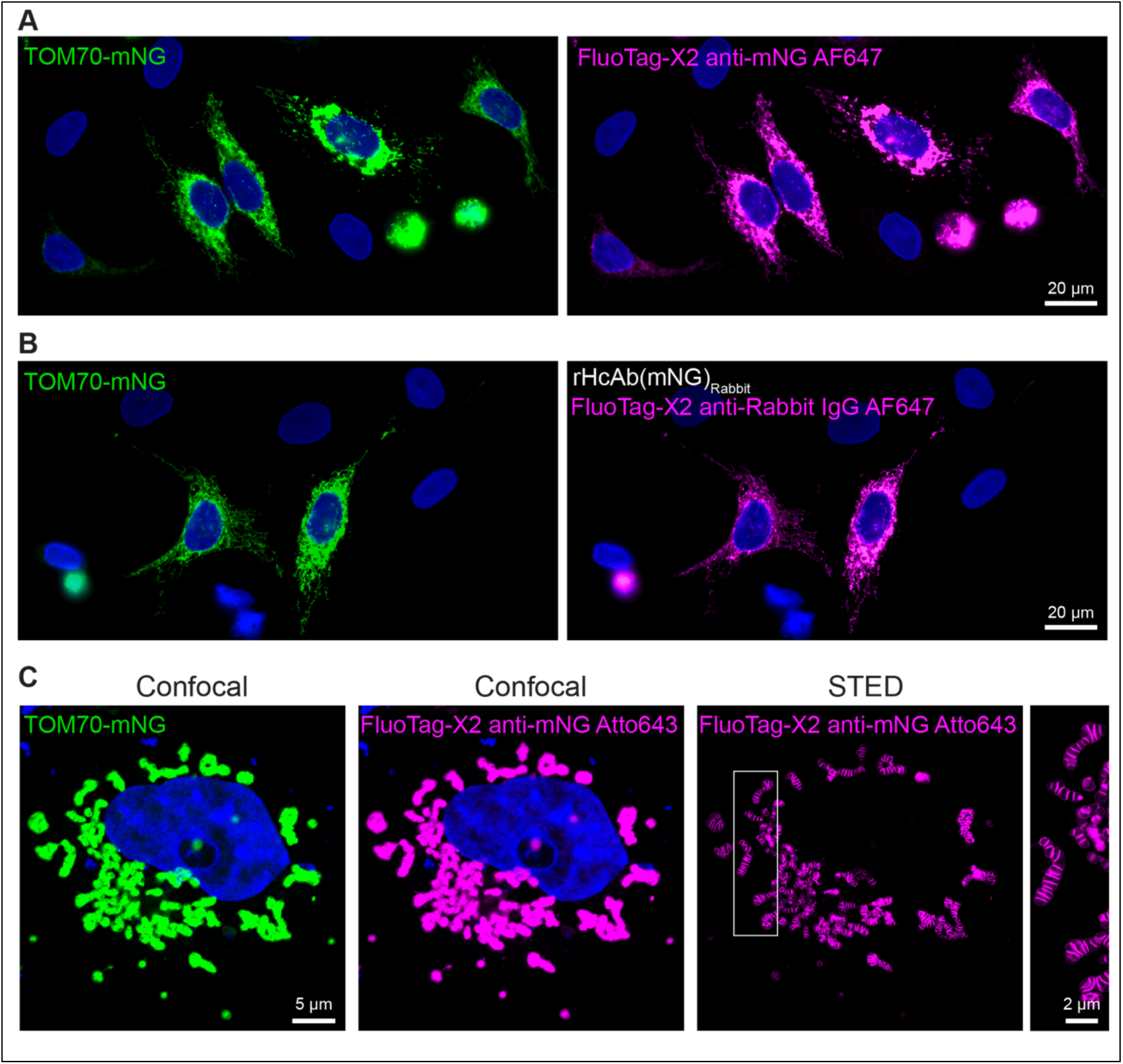
**Immunofluorescence detection of mNeonGreen in PFA-fixed cells**. (**A, B**) HeLa cells transfected TOM70-mNG were fixed, permeabilized and stained for mNG using different sdAb(mNG)-based reagents. Nuclei were revealed by a counterstain against LaminA/C (blue). (**A**) Direct detection of mNG (green) with FluoTag-X2 anti-mNG AF647 (Magenta). (**B**) Indirect detection using a recombinant sdAb(mNG) fused to rabbit IgG Fc domain (recombinant Heavy-chain Antibody; rHcAb(mNG)_Rabbit_) followed by FluoTag-X2 anti-Rabbit IgG AF647. More exemplary direct and indirect stainings are in Supp. Fig. 5. (**C**) COS-7 cells transfected with TOM70-mNG (green), fixed, permeabilized and directly stained using FluoTag-X2 anti-mNG conjugated to Atto643 (magenta) and imaged by confocal and STED microscopy. Nucleus was counterstained with DAPI. Boxed region in the STED panel is displayed larger in the rightmost panel.

To assess the robustness of mNG detection across different sample preparation protocols, HeLa cells expressing TOM70-mNG were analyzed after fixation using four different methods: paraformaldehyde (PFA), methanol, glyoxal, and glutaraldehyde (Supp. Fig. 6). Subsequent immunostaining with FluoTag-X2 anti-mNG revealed clear and specific labeling of mitochondria in all the chemically fixing conditions tested. As expected, glutaraldehyde fixation led to increased nonspecific signal due to autofluorescence induced by the fixative. These results confirm that the reagent is compatible with a broad range of fixation chemistries.

**Figure 6.**
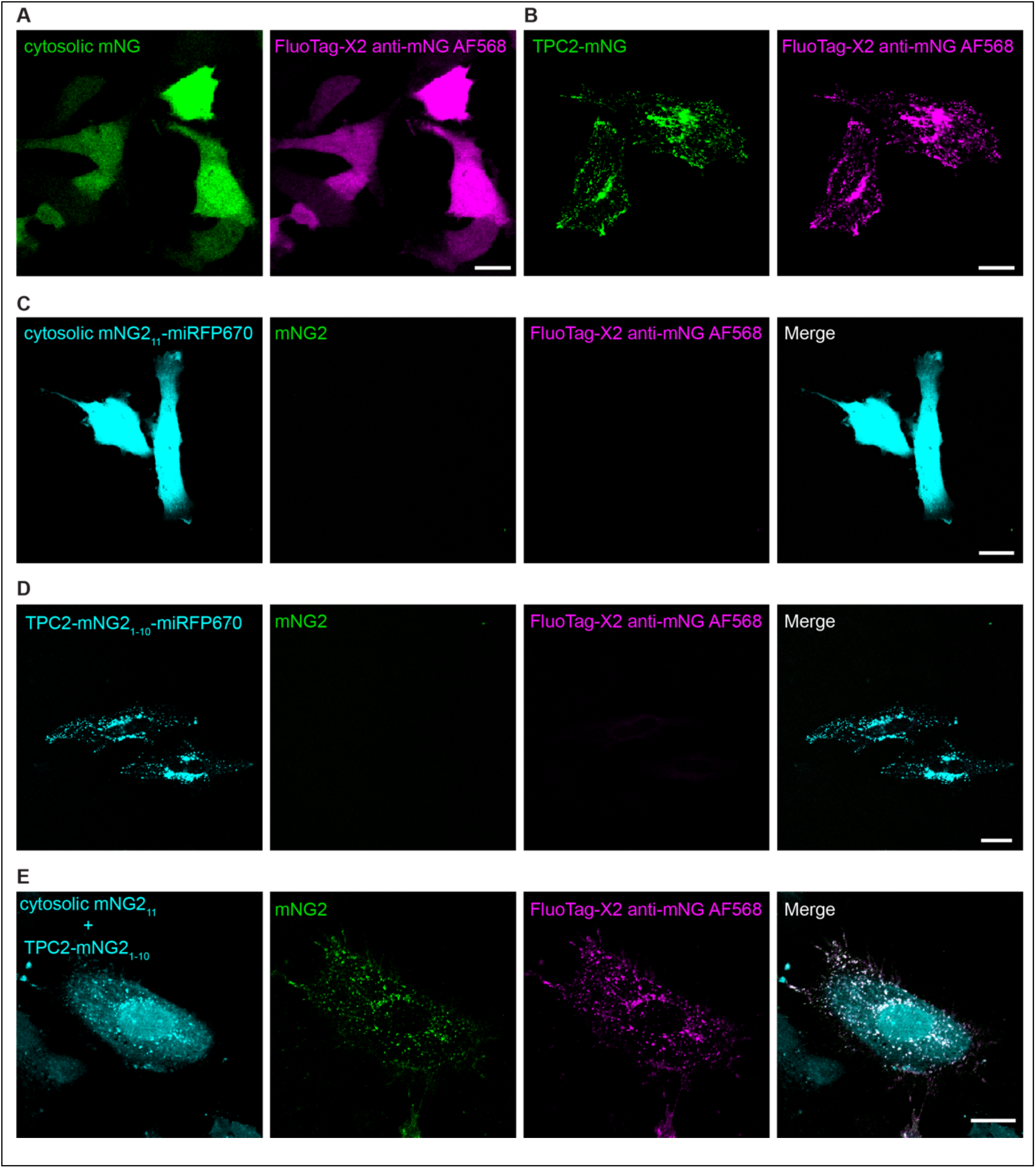
**Binding of sdAb(mNG) to split mNG2**. Both cytosolic mNeonGreen (mNG) (**A**) or lysosomal TPC2 fused to mNG (**B**) are recognized by FluoTag-X2 anti-mNG AF568 when expressed in HeLa cells. Cells expressing cytosolic mNG2_11_ (**C**) or TPC2-mNG_1-10_ (**D**), both fused to miRFP670nano3, show no mNeonGreen fluorescence, and neither fragment is recognized by FluoTag-X2 anti-mNG AF568. (**E**) Co-expression of TPC2-mNG2₁-₁₀-miRFP670nano3 and cytosolic mNG2₁₁-miRFP670nano3 results in reconstitution of mNG2 at lysosomes, evidenced by the recovery of mNeonGreen fluorescence. Recognition by sdAb(mNG) is likewise restored, demonstrating selective binding to the reconstituted fluorophore. Scale bars, 20 µm.

### sdAb(mNG) recognizes only reconstituted split mNG2

We next tested whether sdAb(mNG) recognizes split mNG2^18^, which differs from mNG by five amino acid substitutions (K128M, S142T, R150M, G172V, and K213M). We first assessed recognition of the individual split mNG2 fragments. The small fragment, mNG2₁₁ (β-strand 11 of the barrel, see Fig. 2a), was expressed as a soluble cytosolic protein, whereas the larger fragment, mNG2₁–₁₀ (β-strand 1 to 10, Fig.2A), was fused to the cytosolic C-terminus of lysosome-localized TPC2. Both constructs also contained miRFP670nano3^19^ as a fluorescent reporter for transfection and subcellular localization. As expected, neither split fragment was fluorescent when expressed individually. Surprisingly, FluoTag-X2 anti-mNG AF568 failed to recognize either split fragment (Fig. 6C, D), indicating that the sdAb epitope is generated only upon assembly of the complete mNG2 structure. In contrast, co-expression of both split mNG2 fragments reconstituted mNG2 fluorescence at lysosomes, demonstrating that the sdAb selectively recognizes the fully assembled mNG2 (Fig. 6E).

### Intrabody function of sdAb(mNG) inside living cells

To assess the functionality of sdAb(mNG) in the cytoplasm of living cells, two plasmids encoding a sdAb(mNG) fused to mScarlet-I^20^ fusion or TOM70-mNG, respectively, were transiently transfected in mammalian cells. In the absence of mNG, the sdAb(mNG)-mScarlet fusion was predominantly found in the nucleus (Supp. Fig. 7A), while TOM70-mNG showed the expected localization to mitochondria (Supp. Fig. 7B). Strikingly, when both proteins were co-expressed in the same cell, the sdAb(mNG)-mScarlet-I fusion was re-localized to mitochondria (Fig. 7A). The re-localization efficiency was dependent on the relative expression level of TOM70-mNG with respect to the sdAb(mNG)-mScarlet fusion. At sufficiently high expression of TOM70-mNG, most, if not all, sdAb(mNG)-mScarlet fusion was relocated (Fig. 7A; cell marked with arrowhead) while re-localization was less efficient when TOM70-mNG was expressed at lower levels (Fig. 7A; asterisk). This experiment shows that sdAb(mNG) retained its mNG-binding activity even in the reducing cytoplasmic environment, functioning as an intrabody.

**Figure 7.**
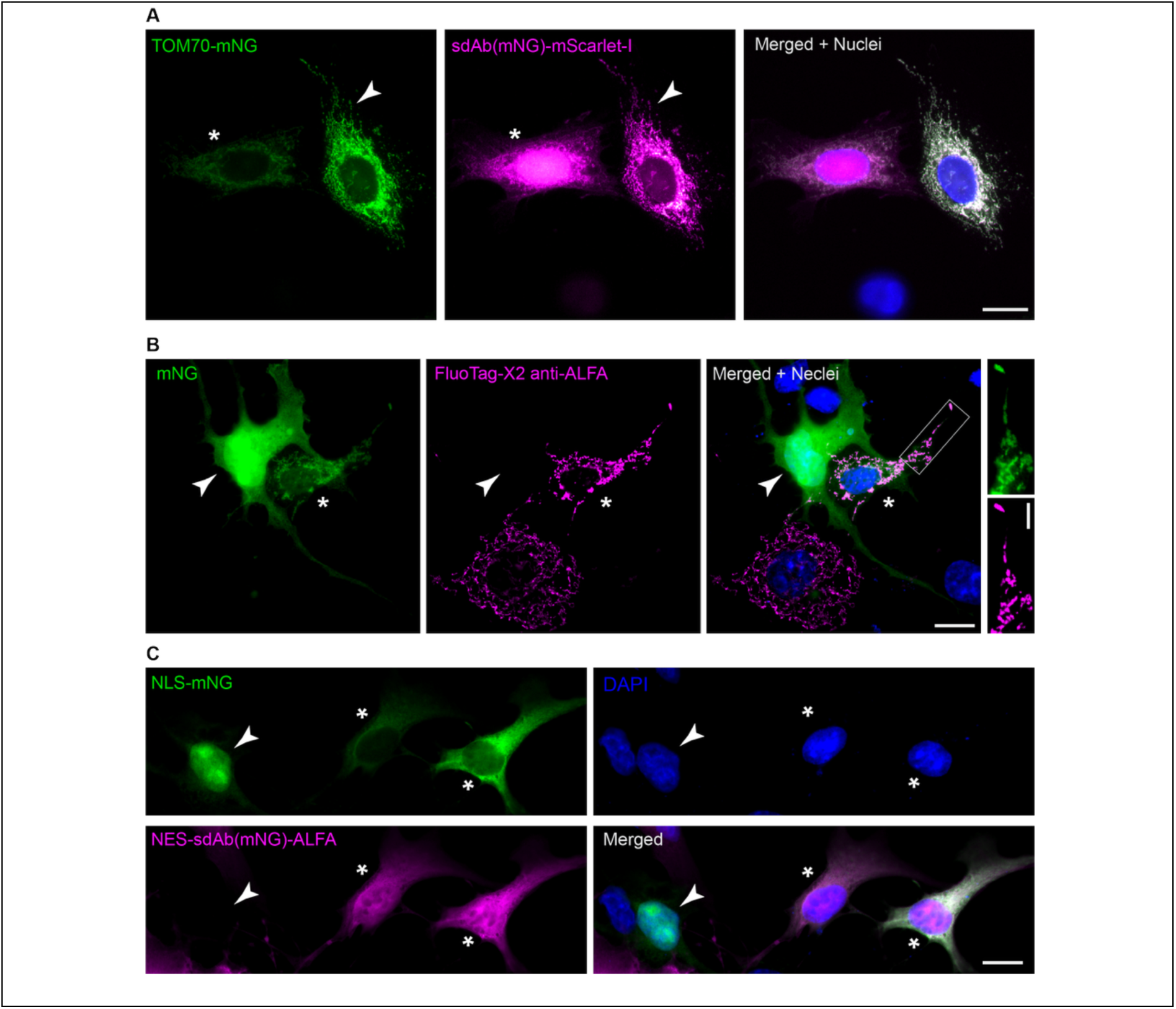
Interaction and re-localization of mNeonGreen using the sdAb anti-mNeonGreen in living cells. **(A)** HeLa cells were co-transfected with plasmids encoding TOM70-mNG (green) and sdAb(mNG)-mScarlet-I (magenta), or each plasmid individually (see Supp. Fig. 7). Cells were counterstained for LaminA/C (blue). In co-transfected cells, sdAb(mNG)-mScarlet-I is re-localized to mitochondria. The arrowhead marks a co-transfected cell with even expression level of both constructs, which results in clear mitochondrial recruitment. In contrast, in cells with non-balanced expression levels (asterisk), the surplus of sdAb(mNG)-mScarlet-I accumulates in the nucleus. Scale bar: 20 µm. **(B)** COS-7 cells were co-transfected with plasmids encoding soluble cytoplasmic mNG and TOM70-sdAb(mNG)-ALFA, which anchors the sdAb to mitochondria. The sdAb(mNG) localization was visualized by staining the ALFA-tag with FluoTag-X2 anti-ALFA AF647 (magenta). In cells expressing mNG alone (arrowhead), mNG is diffusely distributed in the cytoplasm and nucleus. In co-transfected cells (*), soluble mNG accumulates at mitochondria through interaction with mitochondrial sdAb(mNG), as highlighted in the magnified region (rectangle). Scale bars: 20 µm; inset: 5 µm. **(C)** COS-7 cells were co-transfected with NLS-mNG (green) and NES-sdAb(mNG)-ALFA (magenta). SdAb(mNG) localization was detected by staining the ALFA-tag with FluoTag-X2 anti-ALFA AF647 (magenta). Nuclei were counterstained with DAPI (blue). Arrowheads mark cells expressing NLS-mNG alone, where, as expected, the mNG localizes predominantly to the nucleus. Asterisks (*) indicate co-transfected cells in which the NES-bearing sdAb(mNG) which, thanks to the NES, is able to retain most of the mNG in the cytoplasm. Scale bar: 20 µm.

In an inverted system, sdAb(mNG) fused to a mitochondrial targeting sequence (TOM70) efficiently redirected soluble mNG expressed in the cytoplasm of mammalian cells to the mitochondrial surface (Fig. 7B, asterisk), while mNG was more uniformly distributed across the cytoplasm and nucleus in the absence of TOM70-sdAb(mNG) (Fig. 7B, arrowhead). Similarly, mNG, equipped with a nuclear localization signal (NLS), accumulated in the nucleus when expressed alone (Fig. 7C, arrowhead), but was efficiently redirected to the cytoplasm when co-expressed with sdAb(mNG) fused to a nuclear export signal (NES) (Fig. 7C, asterisks). Because sdAb(mNG) is smaller than the passive diffusion cutoff of nuclear pore complexes^21^, excess NES-fused sdAb(mNG) can equilibrate between nucleus and cytoplasm, explaining its presence in both compartments.

We then moved on to test if the sdAb(mNG) is functional as an intrabody when expressed intracellularly in primary rat hippocampal neurons. For this, we used a previously published sdAb anti-Synaptogamin 1 (NbSyt1), which strongly localized to presynaptic terminals and enabled us to track the movement of synaptic vesicles through axons without perturbing neuronal synaptic physiology when expressed as an intrabody^12^. We generated an adeno-associated virus (AAV) encoding for the NbSyt1 fused to mNeonGreen (NbSyt1-mNG), and a second AAV encoding for sdAb(mNG) fused to mScarlet-I (sdAb(mNG)-mScarlet) to co-infect primary neuronal cultures. The sdAb(mNG) clearly works as an intrabody inside neurons, since NbSyt1-mNG strongly co-localizes with sdAb(mNG)-mScarlet not only in static synaptic vesicle clusters at pre-synapses but also when synaptic vesicles or immature packages are moving through the axons (Fig. 8 and Supp. Movie 2 & 3).

**Figure 8.**
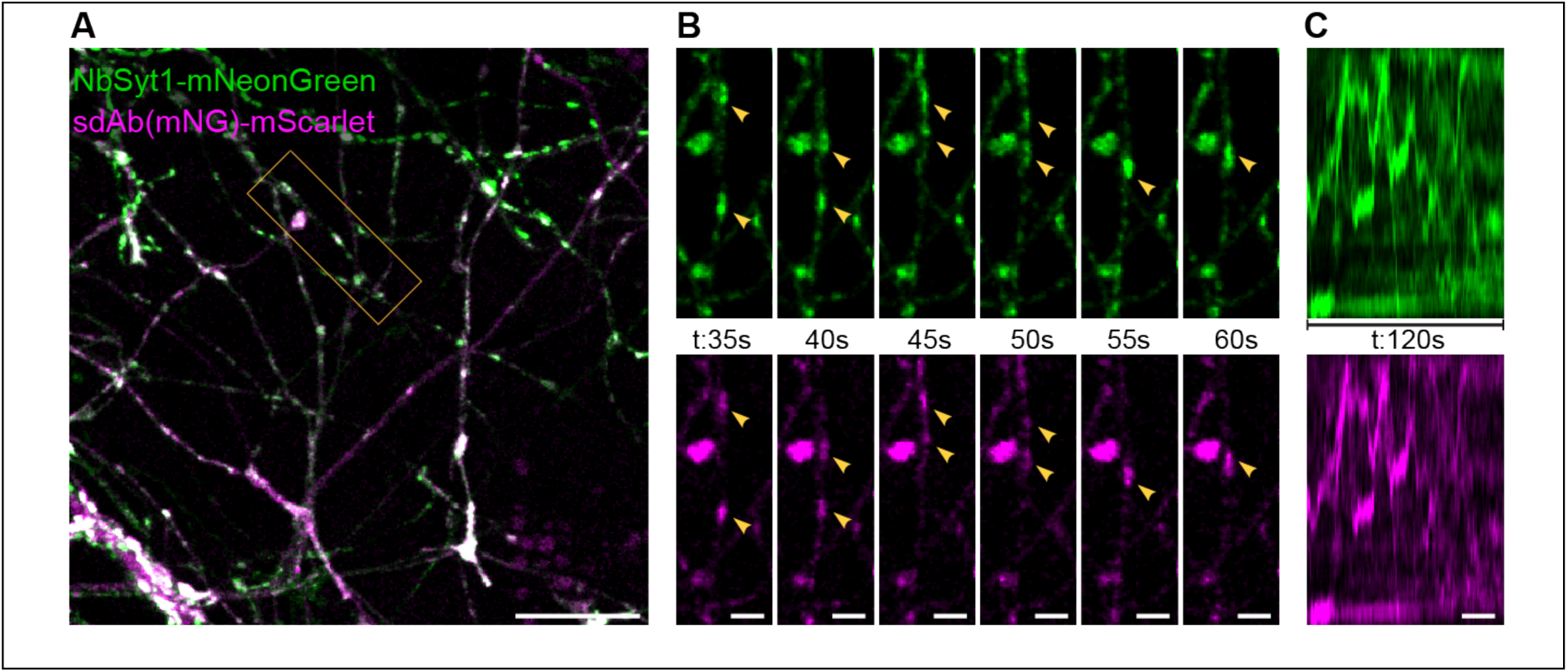
Intracellular expression and function of sdAb(mNG) in hippocampal neurons. (**A**) Overview of rat hippocampal neurons AAV-transduced with the anti NbSyt1-mNG and with the sdAb(mNG) fused to mScarlet-I (sdAb(mNG)-mScarlet). Scale bar: 10 µm (**B**) Selected timeframes of the live time-lapse recording. Yellow arrowheads point to colocalization of mNeonGreen and mScarlet-I in living neurons. Scale bar: 2.65 µm (**C**) Kymographs analysis of the panels shown in B, displaying the movement of both intrabodies in matching directions. Scale bar for the time dimension: 20 s. Note the matching trajectories in both channels, also visible in Supp. Movie 2.

### Targeted degradation of proteins fused to mNG *in vivo* using sdAb(mNG) intrabody

To enable functional interrogation of endogenous Tbxta in zebrafish, we generated a CRISPR/Cas9-mediated knock-in line in which mNeonGreen (mNG) is fused to the C-terminus of Tbxta (Fig. 9A-C). Tbxta (also known as no tail, ntl) is the zebrafish orthologue of the T-box transcription factor Brachyury and plays a central role in mesoderm specification and notochord formation during early development^22,23^. Consistent with this function, the resulting Tbxta–mNG reporter faithfully recapitulates the expected expression pattern in the notochord and tailbud, with clear nuclear localization at single-cell resolution (Fig. 9B-C’). This establishes a robust system for directly visualizing Tbxta dynamics in vivo while preserving endogenous regulation, and provides a defined entry point for intrabody-mediated manipulation of the tagged protein.

**Figure 9.**
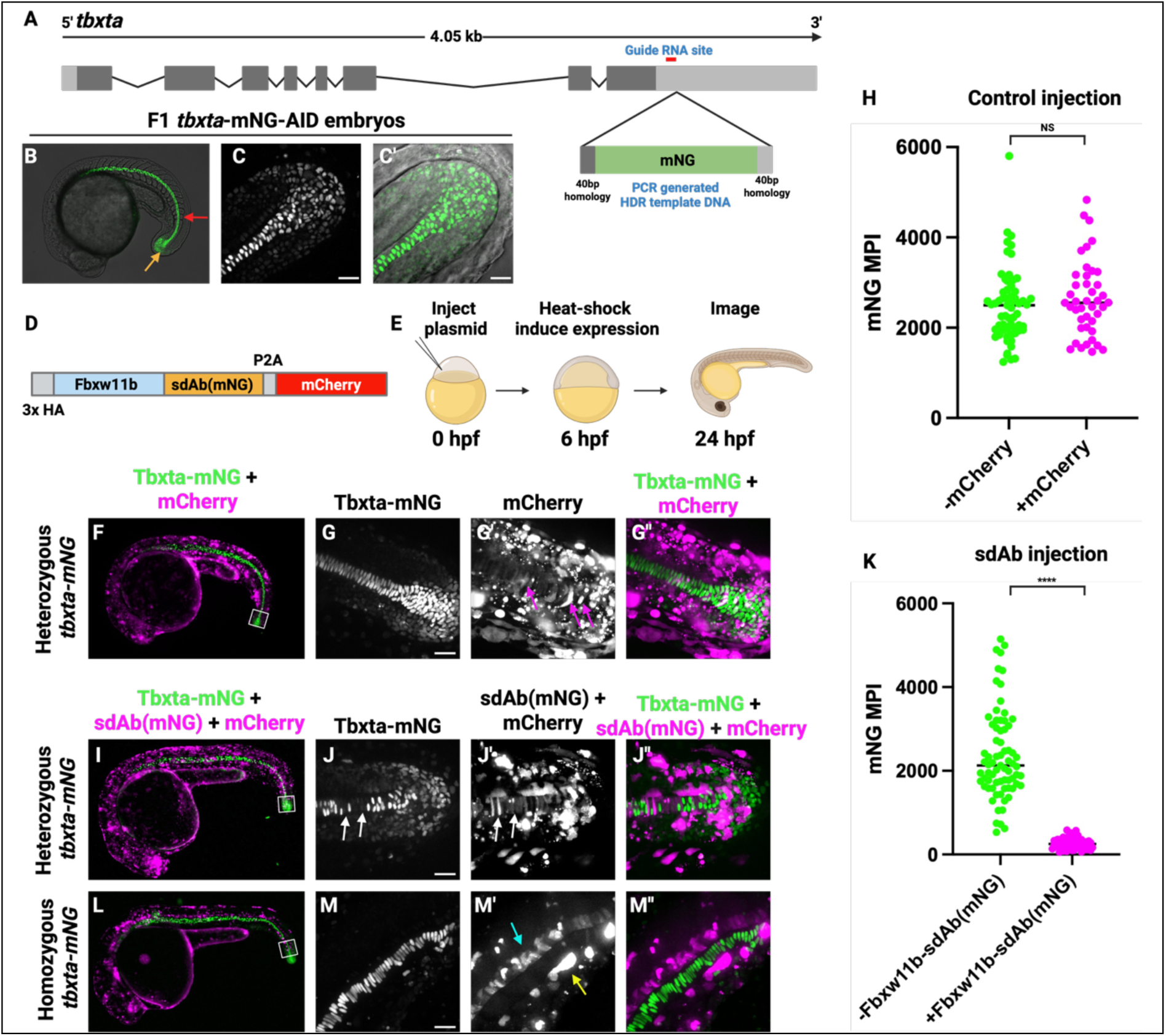
Depletion of mNG tagged endogenous Tbxta from zebrafish embryos using sdAb(mNG). (**A**) Strategy for tagging the endogenous *tbxta* locus with mNG. CRISPR/Cas9 was used to cut the genomic DNA adjacent to the stop codon and a PCR generated repair template with the mNG coding sequence flanked by 40bp of homology was inserted into the cut site. (**B**) Stable F1 embryos exhibit mNG fluorescence in the notochord (red arrow) and tailbud (orange arrow), where *tbxta* is normally expressed. (**C, C’**) High resolution confocal images showing nuclear localization of Tbxta-mNG in the posterior notochord and tailbud without (**C**) and with (**C’**) DIC overlay. (**D**) Construct designed to deplete mNG using the zebrafish F-box protein Fbxw11b fused to the sdAb(mNG) followed by the P2A viral peptide and an mCherry reporter, which was cloned into a heat-shock inducible plasmid. (**E**) Experimental strategy for depleting mNG. The *HS:fbxw11b-sdAb(mNG)-p2a-mCherry* or control *HS:mCherry* plasmids were injected into 1-cell stage embryos, which are inherited mosaically in zebrafish embryos. At 6 hours post fertilization (hpf) embryos were heat-shocked to induce expression and then imaged at 24 hpf for both mNG and mCherry fluorescence. (**F**) A heterozygous *tbxta-mNG* embryo injected with *HS:mCherry* plasmid showing both Tbxta-mNG (green) and mCherry (magenta) fluorescence. (**G-G’’**) The boxed region in F was imaged on a confocal microscope showing Tbxta-mNG (**G**), mCherry (**G’**), or an overlay of both (**G’’**). Notochord cells expressing mCherry (magenta arrows in G’) do not show depletion of Tbxta-mNG (**G’’**). (**H**) Quantification of mNG fluorescence mean pixel intensity in nuclei of notochord cells with and without mCherry shows there is no significant difference between the two (N=5 embryos, 61 cells without mCherry, 42 cells with mCherry). (I) A heterozygous *tbxta-mNG* embryo injected with *HS:fbxw11b-sdAb(mNG)-p2a-mCherry* plasmid showing both Tbxta-mNG (green) and mCherry (magenta) fluorescence. (**J-J’’**) The boxed region in I was imaged on a confocal microscope showing Tbxta-mNG (**J**), mCherry (**J’**), or an overlay of both (**J’’**). (**J**) Gaps in notochord Tbxta-mNG fluorescence (white arrows) correspond to cells that have Fbxw11b-sdAb(mNG) as indicated by mCherry fluorescence (white arrows in J’). (**K**) Quantification of mNG fluorescence mean pixel intensity in nuclei of notochord cells with and without Fbxw11b-sdAb(mNG) shows there is a significant reduction in Tbxta-mNG fluorescence in Fbxw11b-sdAb(mNG) containing cells (N=5 embryos, 73 cells without Fbxw11b-sdAb(mNG), 51 cells with Fbxw11b-sdAb(mNG). (**L**) A homozygous tbxta-mNG embryo injected with HS:fbxw11b-sdAb(mNG)-p2a-mCherry plasmid showing both Tbxta-mNG (green) and mCherry (magenta) fluorescence. (**M-M’’**) The boxed region in L was imaged on a confocal microscope showing Tbxta-mNG (**M**), mCherry (M’), or an overlay of both (**M’’**). Tbxta is essential for cells to adopt a notochord fate. Cells containing Fbxw11b-sdAb(mNG) adopt an alternative floor plate (M’, cyan arrow) or hypochord fate (M’, yellow arrow), with no mCherry fluorescence observed in the notochord (M’, M’’) (N=3 homozygous embryos). Scale bars = 20 μm. This entire Figure was created with the help of https://BioRender.com

We exploited this system to achieve targeted depletion of Tbxta–mNG using a degradation module consisting of the zebrafish F-box protein Fbxw11b fused to the sdAb(mNG) (Fig. 9D-E), based on methods developed for targeted degradation of GFP-tagged proteins^24^. Mosaic expression of this construct, induced via heat shock, led to a striking and cell-autonomous loss of Tbxta–mNG specifically in Fbxw11b-sdAb(mNG)-positive cells identified by the mCherry reporter fluorescence (Fig. 9I-J’’), whereas control injections had no effect on fluorescence levels (Fig. 9F-H). Quantitative analysis confirmed a significant reduction of nuclear mNG signal upon nanobody-mediated targeting (Fig. 9K). In homozygous embryos, depletion of Tbxta resulted in a clear cell fate switch, with affected cells failing to contribute to the notochord and instead adopting alternative identities such as floor plate or hypochord (Fig. 9L-M’’). Together, these results demonstrate that sdAb(mNG) as an intrabody can be harnessed to efficiently and spatially resolve the degradation of endogenous proteins *in vivo*, providing a powerful and versatile strategy for dissecting protein function during development and beyond.

## Discussion

The here characterized sdAb(mNG) is a highly effective binder of mNeonGreen. Its development was motivated by the widespread use of mNeonGreen as a protein tag, owing to its exceptional brightness, rapid chromophore maturation, strict monomericity, and robust photostability^1^. Despite these advantages, the repertoire of affinity reagents for mNG-tagged proteins has remained limited compared with that available for other fluorescent proteins. sdAb(mNG) fills this gap by providing a high-affinity binder that enables applications not readily supported by conventional anti-mNG binders.

The structural and biophysical data together provide a coherent mechanistic account of the reagent’s remarkable combination of high affinity and broad resilience. The crystal structure reveals engagement of all three CDR loops against the β-barrel lateral surface through a chemically diverse interface. It is this interaction richness that underpins the sub-nanomolar affinity and slow dissociation rate. The same features explain why the complex survives high-salt buffers, reducing agents, detergents, and acidic conditions down to pH 2.2.

An underappreciated consequence of tight, structurally rich binding is thermal stabilization of the target protein. The exceptional thermostability and refolding capacity of sdAb(mNG) itself, surviving high temperatures and recovering function after exposure to guanidinium hydrochloride, SDS, or DTT, stand out among affinity reagents and have immediate practical value for affinity resin regeneration, long-term storage, and repeated-use workflows, including large-scale affinity enrichment coupled to mass spectrometry proteomics.

The specificity analysis confirmed clean discrimination against a panel of structurally related FPs, attributable to divergence at contact residues. More mechanistically revealing is the behavior toward split mNG2: sdAb(mNG), which recognizes the reconstituted split mNG2 complex but fails to bind the isolated mNG2_1-10_ fragment, even though the epitope lies outside the sequence corresponding to the missing eleventh β-strand. This implies that the large fragment, without its terminal strand, adopts a conformation sufficiently different from the native barrel to be unrecognized, adding to growing evidence that deletion of terminal strands perturbs the global fold of β-barrel fluorescent proteins beyond the local deletion site^18,25,26^. From a practical standpoint, selective recognition of fully reconstituted split mNG2, but not of its components, turns sdAb(mNG) into a coincidence detector for protein–protein interactions reported by the split mNG2 system, enabling affinity purification or in situ detection of interacting complexes.

The reduction in linkage error with a directly conjugated sdAb is a highly significant feature in super-resolution modalities^7,27,28^, where the separation between fluorophore and target competes with effective resolution, as exemplified by the resolved mitochondrial outer membrane architecture. Compatibility with chemically diverse fixation protocols, PFA, methanol, glyoxal, and glutaraldehyde, extends the reagent’s utility across laboratories with divergent sample preparation traditions. Looking ahead, the defined and geometrically favorable C-terminal conjugation makes sdAb(mNG) an attractive candidate for correlative light and electron microscopy (CLEM) and cryo-EM fiducial strategies^29,30^, and expansion microscopy workflows^31^, where the compact size of single-domain antibodies is a structural asset.

Perhaps the most demanding test of any sdAb binder is whether it retains function in the cytoplasm of living cells^11^. The reducing intracellular milieu disfavors disulfide bond formation and can destabilize antibody-derived domains that depend on it. For this reason, many sdAbs that perform well in biochemical assays fail as intrabodies. Relocalization assays show that sdAb(mNG) retains full activity in the cytoplasm, thereby validating it as an intrabody. In primary rat hippocampal neurons, it enables functional co-tracking of synaptic vesicles without disrupting synaptic function, demonstrating robustness and functional orthogonality in differentiated cells. This facilitates multiplexed intrabody labeling for live vesicle imaging, which has previously been constrained by the limited availability of suitable reagents.

In our zebrafish proof-of-concept application, *in vivo* deployment of sdAb(mNG) leverages all its key features: tight binding, intracellular function, and tolerability of fusion moieties. Using sdAb(mNG) in an Fbxw11b-based degradation system, heat-shock-induced mosaic expression caused cell-autonomous depletion of endogenous tagged Tbxta-mNG, leading to a cell-fate switch aligned with Tbxta’s role in mesoderm specification. This experiment shows that any protein tagged with mNG in organisms with conditional expression can be targeted for rapid, localized degradation or functional interference. Coupled with the expansion of CRISPR-mediated mNG knock-in lines, this approach enables precise analysis of protein function during embryonic development, organ formation, and tissue maintenance, especially where traditional genetic methods elicit compensatory responses or lack spatial resolution.

The development of sdAb(mNG) highlights a broader challenge in the life sciences: experimental reproducibility in antibody-reagent-dependent assays. Polyclonal antisera, used for decades, are inherently irreproducible once the immunized animal is exhausted; batch variability in titer, specificity, and background is well documented, and the source animal’s death marks the reagent’s end^32^. This affects scientific literature: many commercial antibodies fail to recognize their targets, and published results can’t be reproduced when the original stocks are depleted. Recombinant, sequence-defined probes like sdAb(mNG) are the opposite: their binding sequences are fully known and can be produced indefinitely from a plasmid in standardized systems. Labs can, in principle, produce identical reagents, validate against the same epitope, and compare results with those published. This reproducibility-by-design advocates for replacing serum-derived reagents with recombinant, well-characterized binders across the life sciences, a transition made more feasible by tools like this.

With mNG as one of the leading fluorescent protein tags, sdAb(mNG) provides a foundation for expanding the range of tools across structural, synthetic, developmental, and proteomic research, limited only by community creativity.

## Data Availability

The crystal structure is deposited on the RCSB Protein Data Bank under the code 30TF. Raw X-ray diffraction and SAXS data will be made available upon publication via Zenodo. Other data and materials can be made available upon reasonable request.

## Acknowledgment

We thank Jannik Hentze for great technical support. F.O. was supported by Deutsche Forschungsgemeinschaft (DFG) through the SFB 1286 (project Z04). P.K. was supported by the Research Council of Norway grants BIOPROM (grant number 324877) and NeuroConvergence (grant number 331725). We acknowledge the Core Facility for Biophysics, Structural Biology, and Screening (BiSS) at the University of Bergen, which has received infrastructure funding from the Research Council of Norway through NORCRYST (grant number 245828) and NOR–OPENSCREEN (grant number 245922). V.K. has been supported by the European Union’s Horizon 2020 research and innovation program under the Marie Skłodowska-Curie grant agreement 101151099. We further wish to acknowledge experimental beamtime and beamline support at DESY synchrotron beamlines P11 and P12.

## Methods

### Alpaca Immunizations

All animal work was conducted in accordance with ethical regulations for animal research and testing. The experiments did not require formal ethical approval but were reported to and approved by the local authorities (LAVES, Niedersachsen, Germany). Two alpacas were immunized six times at 14- to 42-day intervals, with a total of 0.5 mg of mNeonGreen. GERBU FAMA was used as an adjuvant for all immunizations. Five days after the final immunization, 100 mL of peripheral blood was collected from each animal.

### Preparation of Phagemid Libraries

A phagemid library was constructed to select mNeonGreen-specific binders. The library was generated from total ribonucleic acid (RNA) isolated from peripheral blood mononuclear cells (PBMCs) obtained from fresh alpaca blood. PBMCs were isolated using Ficoll-Paque PLUS (GE Healthcare, Uppsala, Sweden), and total RNA was extracted using a NucleoSpin RNA Plus kit (Macherey-Nagel, Düren, Germany). The isolated RNA was used for reverse transcription with SuperScript IV Reverse Transcriptase (Thermo Fisher Scientific, Waltham, MA, USA). SdAb-encoding sequences were amplified by a two-step nested polymerase chain reaction (PCR) using primers CALL01 and CALL02, followed by primers F1 and R1 in the second PCR^33^. The final PCR products were cloned into a pHen2-derived phagemid vector and transformed into TG1 cells, yielding sdAb libraries with a complexity of >5 × 10⁸ individual clones.

### Phage Display

A total of three consecutive biopanning rounds were performed using mNG immobilized on magnetic beads. In each panning round, phages were incubated for 1 h at room temperature with mNG-charged beads pre-blocked with 20 µg/mL bovine serum albumin (BSA). Bound phages were proteolytically eluted and used to infect *E. coli* TG1 cells. Infected cells were grown overnight at 37 °C. The following day, the culture was infected with an M13KO7 helper phage. For phage production, cultures were grown in the presence of appropriate antibiotics over night at 30 °C. Phages were purified from cleared culture supernatants by repeated precipitation with PEG-8000.

### Protein expression and purification

Recombinant proteins used in this study were expressed in *Escherichia coli* under the control of an IPTG-inducible promoter from expression vectors carrying a ColE1 origin and kanamycin resistance. All proteins contained an N-terminal His-tag. For expression, *E. coli* cultures were grown in Terrific Broth (TB) and induced with 0.5 mM IPTG, followed by overnight expression at 23 °C. Cells were harvested by centrifugation, resuspended in LS buffer (50 mM Tris/HCl pH 7.5, 300 mM NaCl) supplemented with 15 mM imidazole and 10 mM DTT and lysed by sonication. Lysates were cleared by centrifugation and purified by IMAC chromatography. For applications requiring higher purity, the His-tag was proteolytically removed before further purification by size-exclusion chromatography (Superdex 75 or Superdex 200 (Cytiva)). Protein purity was assessed by SDS-PAGE.

### Affinity capture from *E. coli* lysate

Affinity capture experiments from *E. coli* lysate were performed using a defined amount of affinity-purified recombinant mNeonGreen. For each experiment, 20 µL of mNeonGreen Selector resin (NanoTag Biotechnologies; Cat. No. N3210) was incubated with 1 mL of lysate containing 3 µM mNeonGreen for 1 h at room temperature. After binding, the resin was washed three times in batch with 1 mL of PBS, transferred to a MiniSpin column, and washed twice with 0.6 mL of PBS. For elution, the resin was resuspended in SDS sample buffer and heated to 95 °C for 5 min. As a specificity control, mNeonGreen Selector resin was treated in parallel using *E. coli* lysate lacking mNeonGreen. Samples were resolved by SDS-PAGE and analyzed by Coomassie staining.

### Biolayer Interferometry (BLI)

Binding kinetics between sdAb(mNG) and mNeonGreen were determined by biolayer interferometry (BLI) using an Octet^®^ R8 instrument (Sartorius) at 22 °C with continuous shaking at 1000 rpm. Measurements were performed using Octet^®^ High Precision Streptavidin (SAX) biosensors. Site-specifically biotinylated sdAb(mNG) (clone 1E2) was immobilized onto SAX biosensors at a concentration of 0.6 µg/mL for 200 s following an initial baseline step (60 s) in kinetics buffer (PBS supplemented with 0.01% BSA and 0.002% Tween-20). Remaining streptavidin binding sites were blocked using 100 µg/mL biotin for 120 s. Sensors were subsequently equilibrated in the kinetics buffer for 120 s prior to association measurements. Association kinetics were recorded by exposing the sensors to soluble mNeonGreen in a two-fold dilution series ranging from 20 nM to 0.3125 nM for 900 s. Dissociation was monitored for 2700 s in the kinetics buffer. Sensorgrams were reference-subtracted and processed using inter-step correction and Savitzky–Golay filtering. Kinetic parameters were obtained by global fitting using a 1:1 Langmuir binding model. Association (k_on_) and dissociation (k_off_) rate constants were determined from the fits, and the equilibrium dissociation constant (K_D_) was calculated as k_off_/k_on_.

### Thermostability assays

To assess the thermostability of non-complexed mNeonGreen, 50 µL of a 25 µM mNeonGreen solution in PBS was incubated in a PCR thermocycler for 15 min at defined temperatures ranging from 63 °C to 77 °C. Following incubation, samples were cooled to room temperature and centrifuged (1 min, 500 rcf). Fluorescence images were acquired under UV illumination. For direct comparison, an identical assay was performed in parallel using a 20% slurry of mNeonGreen Selector preloaded with 50 µM mNeonGreen (concentration refers to resin volume). To evaluate refolding after thermal denaturation in the presence of various reagents, 30 µL of mNeonGreen Selector in PBS was heated to 95 °C in PBS alone or in PBS supplemented with 100 mM DTT, 2 M guanidine hydrochloride (GuaHCl), or 0.5% SDS. After cooling to room temperature, the resin was washed twice with PBS and incubated with 50 µM mNeonGreen for 5 min (concentration is based on the resin volume). Fluorescence images of the settled incubation mixtures were acquired under UV illumination.

### Specificity assays

For equilibrium binding assays, a 40 µL volume of mNeonGreen Selector was washed and incubated in TBS supplemented with 2.5 µM of the indicated recombinantly expressed and affinity-purified fluorescent protein for 1 h at room temperature. After incubation, the beads were sedimented, and a picture was acquired under UV illumination using a Nikon D700 camera equipped with a 105 mm macro lens (Nikon, Minato, Japan).

For ELISA-based specificity assays, microtiter plates were coated overnight at 4 °C with 100 fmol of fluorescent target protein per well. Wells were blocked with CANDOR SmartBlock for 1 h at RT, then washed 3 times with TBS-T. Wells were further incubated with 100 µL of a 3 nM solution of biotinylated sdAb(mNG) for 1 h at RT. After washing three times with TBS-T, Streptavidin–HRP conjugate was applied at a 1:10,000 dilution and incubated for 1 h at room temperature, followed by three additional washes with TBS-T. For detection, 100 µL of TMB substrate (SeramunBlau slow2 50) was added to each well, and the plate was incubated for 5 minutes. The enzymatic reaction was stopped by adding 100 µL of 1 M sulfuric acid. Absorbance was then measured using a microplate reader at 450 nm and 620 nm, respectively. Corrected values were obtained by calculating as (OD_450nm_ – OD_620nm_). All measurements were performed in triplicate.

### Resistance to Stringent Washing and pH before and after binding

A 40 µL volume of mNeonGreen Selector resin (NanoTag Biotechnology; Cat. No. N3210) was washed and saturated with recombinantly expressed, affinity-purified mNeonGreen. Protein binding was performed in Tris-buffered saline (TBS: 25 mM Tris/HCl, pH 7.5; 150 mM NaCl; 2 mM EDTA) or TBS supplemented with the indicated solutions or buffers. The beads were then washed four times with TBS. To test resistance to stringent washing and pH, resins saturated in the absence of additional solutions or buffers were subsequently incubated in 1 mL of the indicated solution or buffer for 2 h at room temperature with shaking. After incubation, the beads were sedimented by centrifugation and imaged under UV illumination using a Nikon D700 camera equipped with a 105 mm macro lens (Nikon, Japan).

### Cell Culture and Transient Transfection

HeLa and COS-7 cells were cultured in DMEM supplemented with high-glucose and stable Glutamine, 10% fetal bovine serum, and 1% penicillin/Streptomycin. Cells were cultured in a humified incubator at 37 °C with 5% CO_2_. For interaction and re-localization studies, cells were transiently transfected with 0.75 µg of plasmid per well of a 12-well plate (as listed in Supplementary Table 2) using the PolyJet transfection kit (SignaGen) according to the manufacturer’s recommendations. COS-7 and HeLa cells were seeded on 12-well plates with cover glass 24 h prior to transfection. Cells were incubated with transfection reagents for 24–48 h. For co-expression experiments, plasmid DNA was premixed in a 1:1 ratio and further processed as described above.

For assessment of split versions of mNG2, HeLa cells were trypsinized and seeded onto CellView slides (Greiner Bio-One). One day after subculturing, cells were transiently transfected with the JetPEI reagent (Sartorius) at a 5:2 ratio with various construct DNA (see Supplementary Table 2). Per well, cells were transfected with 100 ng of plasmid DNA for 4-6 h before the medium was replaced.

### Primary hippocampal neuronal cultures

Wild-type Wistar rats (*Rattus norvegicus*) were obtained from the University Medical Center Göttingen and handled in accordance with the specifications of the University of Göttingen and the local authority, the State of Lower Saxony (Landesamt für Verbraucherschutz, LAVES, Braunschweig, Germany). The local authority approved animal experiments, the Lower Saxony State Office for Consumer Protection and Food Safety (Niedersächsisches Landesamt für Verbraucherschutz und Lebensmittelsicherheit). Rat primary hippocampal neuron cultures for imaging were prepared with minor modifications from^12,34^. Briefly, the brains of P0-P1 rat pups were extracted and placed in cold HBSS. The hippocampi were extracted and placed in a solution containing 10 mL Dulbecco’s Modified Eagle Medium (DMEM), 1.6 mM cysteine, 1 mM CaCl_2_, 0.5 mM EDTA, 25 units of papain (Worthington, #LS003126) per mL of solution, with CO_2_ bubbling, at 37 °C for 1 h. Following enzymatic digestion, the solution was replaced with 10% Fetal Calf Serum in DMEM for 15 min. Subsequently, hippocampi were triturated using a 10 mL pipette in complete Neurobasal-A medium supplemented with 2% B27 and 1% GlutaMax (Thermo Fisher Scientific). Neurons were plated on a glass-bottom 96-well plate, previously coated with poly-L-lysine hydrochloride (PLL, 1 mg/mL) at a density of 5,000 cells per well. After 2 h, the plating medium was replaced with 150 μL of complete neurobasal medium, and neurons were incubated at 37 °C and 5% CO_2_ in a humidified incubator.

### Adeno-Associated Viruses (AAVs)

The sdAb(mNG) was PCR-amplified from the plasmid pNT4316 (see Supp. Table 2) to introduce AgeI and SbfI. The acceptor pAAV vector with the synapsin promoter^12,34^ was digested, dephosphorylated with FastAP, and purified. Both insert and vector were ligated using T4 ligase, then transformed into SURE cells on ampicillin plates. Plasmids from various colonies were sequenced to confirm ITR preservation and sequence accuracy, thereby generating the sdAb(mNG)-mScarlet_AAV. AAVs were generated as previously described^12,34^. Briefly, HEK293 cells were co-transfected with pHelper plasmids (pFΔ6, pRV1, p21; AAV 1/2) and pAAV target plasmid in a 4:1:1:2 molar ratio by use of Lipofectamine 2000 (ThermoFisher Scientific). 72 h post-transfection, cells were harvested and lysed by 4 cycles of freezing and thawing, followed by treatment with Benzonase nuclease. After pelleting cellular debris (16.9 *g* rcf^−1^, 30 min at 4 °C), supernatants were filtered, aliquoted, and snap-frozen in liquid nitrogen before being stored at-80 °C until use.

### Cell Infection

Rat hippocampal neurons (DIV ∼ 7-11) were co-transduced with AAVs containing the sequence for expression of directly fluorescently labeled intrabodies: NbSyt1-mNG^12^) and sdAb(mNG)-mScarlet. Aliquots of AAVs were thawed on ice and diluted in conditioned neuronal medium; serial dilutions were performed to obtain final viral concentrations ranging from 1:100 to 1:1600. The same conditions were applied to both same-day co-infection experiments and to delayed infection experiments, in which the second virus (sdAb(mNG)-mScarlet) was added 72 h prior to imaging (DIV11).

### Immunostainings

Transiently transfected COS-7 and HeLa cells were fixed with 4% (w/v) paraformaldehyde (PFA) in PBS (pH ∼7.5) for 30 min at room temperature (RT) with gentle rocking. Residual aldehydes were quenched with 0.1 M NH₄Cl in PBS for 20 min. Cells were then blocked and permeabilized in PBS containing 10% normal goat serum and 0.1% Triton X-100 for 30 min.

Detection of mNG was performed with the sdAb(mNG) and derivatives described in this work (FluoTag®-X2 anti-mNeonGreen (Cat. No. N3202), Fc fusions: rHcAb(mNG)_Mouse_; rHcAb(mNG)_Rabbit_ (Cat. Nos. N3282 and N3283, respectively), also detailed in Supplementary Table 3. Other affinity reagents from NanoTag Biotechnologies GmbH were FluoTag^®^-X2 anti-ALFA^®^- Alexa Fluor™ 647 (Cat. No: N1502); FluoTag^®^-X2 anti-Rabbit-IgG Alexa Fluor™ 647 (Cat. No: N2402); FluoTag^®^-X2 anti-Mouse IgG1 Alexa Fluor™ 647 (Cat. No: N2002); FluoTag^®^-X2 anti-Mouse IgG2 Alexa Fluor™ 568 (Cat. No: N2702); from Abcam, the anti-Lamin (Mouse IgG2; ab238303). A full list of antibodies with more details can be found in Supplementary Table 3. For Supplementary Figure 6, transfected HeLa cells were fixed using different methods. Fixation with 4% PFA was performed as described above. Alternatively, cells were treated with ice-cold 100% methanol for 15 min, 3% glyoxal solution (in 19.9% ethanol and 0.75% acetic acid) for 30 min, or 2% glutaraldehyde (v/v) in 0.1 M phosphate buffer for 1 h at RT. Quenching, blocking, and permeabilization were performed as described for PFA fixation. Samples were mounted in Mowiol for imaging.

For assessment of split versions of mNG2, 24 hours after transient transfection, HeLa cells were fixed with 4% PFA (in PBS); note that the fluorescent holo-proteins retain fluorescence after fixation (i.e., mNG, reconstituted mNG2, and miRFP670nano3). Cells were permeabilized and blocked with 0.1% Triton X-100 and 10% natural goat serum in PBS for 15 min and incubated with a 1:500 dilution (5 nM final concentration) of FluoTag-X2 anti-mNG Alexa Fluor™ 568 (Cat. No: N3202) for 1 h at RT with gentle agitation. Cells were washed 3 times for 5 min with PBS.

### Microscopy

Immunofluorescence images presented in Figs. 5, 7, and Supp Figs. 5–7 were acquired using a Zeiss Axio Imager Z1 microscope. Image acquisition was performed with a 63× EC Plan-Neofluar 1.25 oil immersion objective, and images were processed using ZEN software (version 2). Confocal and STED images in Figure 5 were acquired via an inverted Abberior STED Expert line microscope (Abberior Instruments, Göttingen, Germany) equipped with a UPLXAPO 100×1.4 NA oil immersion objective (Olympus). The excitation lasers at 488 nm and 640 nm were set to 1 μW, and the 775 nm depletion laser was set to 5 mW, as measured at the sample. The imaging settings were as follows: pixel size of 20 nm, dwell time of 10 μs, and signal accumulation over 2 lines. Both confocal and STED images were collected from the 640 nm channel, whereas only confocal images were acquired from the 488 nm channel. For Figure 6 (split mNG), cells were imaged using a Nikon A1R laser-scanning confocal equipped with a Plan Fluor 40x oil DIC H N2 (NA: 1.3) objective, in Channel-Series mode. Green, red, or far-red fluorophores were alternately excited (ex/em): 488/525 nm, 561/595 nm, and 640/700 nm, respectively.

Live imaging on rat primary hippocampus neurons (Fig. 8) was performed using a Stellaris 8 confocal microscope (Leica Microsystem, Germany) equipped with a Plan Apochromat 100×, 1.4 NA oil immersion objective (Immersion Oil Type F, 11944399, Leica Microsystem), HyDs Sensors, an Oko Touch Environmental control, and LAS X software (v4.9.0.30221). The White-Light Laser (WLL) was equipped and tuned to 504 nm and 570 nm, and the HydS sensor range was adjusted to optimize yield while minimizing fluorophore crosstalk. Excitation laser power for both channels was kept below ∼10% to minimize photobleaching and phototoxicity. Time-lapse movies were acquired frame-by-frame (scan speed of 400 Hz; pixel dwell time of 2.0625 µs) at an imaging rate of 0.77 frames per second for 120 frames. Images were collected at 512 × 512 pixels with a zoom factor of 2.5, corresponding to a pixel size of ∼90 nm. For Fig. 9, zebrafish embryos were mounted in 1% low-melt agarose containing tricaine for imaging. They were first imaged on a Leica DMI6000b inverted compound microscope with a 5x objective, then transferred to a spinning-disk confocal microscope (Nobska Imaging, Inc.) for high-resolution imaging. This microscope consists of a Zeiss Axio Imager A2 frame, a Borealis-modified Yokogawa CSU-10 spinning-disk, an ASI 150-micron piezo stage controlled by an MS2000, an ASI filter wheel, and a Hamamatsu ImagEM X2 EM-CCD camera. Images were collected using a W Plan-Apochromat 40x/1.0 NA water immersion objective.

### Image analysis and figure generation

All images were adjusted and adjusted using Fiji (ImageJ, v2.16.0/1.54p). In Fig. 8, the two channels were split and processed individually. Image processing was performed using Fiji (ImageJ, v2.16.0/1.54p); for both channels, photobleaching correction was first applied (exponential fit, v2.1.0, Miura K., 2020), followed by background subtraction (rolling ball radius = 25 px) and median filtering (radius = 0.25 pixel). For visualization purposes, lookup tables were adjusted without affecting raw intensity values. Regions of interest containing moving vesicular structures were isolated and used to generate kymographs for each fluorescence channel. Dendrites in the region of interest designated to be further analyzed were manually tracked through the segmented line tool based on the background fluorescence coming from the structure. Kymographs for each channel were plotted using the Multi Kymograph plug-in (linewidth = 9 pixels). For Fig. 9 mNG confocal image quantification, rolling-ball background subtraction (radius = 50) was performed in Fiji. The freehand selection tool was used to select the nuclear region and determine the mean gray value.

### Crystal structure determination

mNG and sdAb(mNG) were mixed at a 1:1.2 molar ratio and incubated on ice for 30 min. The complex was concentrated at 4000 rpm using a 10-kDa cut-off Amicon centrifugal filter unit and then injected onto a Superdex 200 Increase column equilibrated in phosphate-buffered saline (PBS). The main peak fractions were concentrated as above to 10–15 mg/ml and used to prepare crystallization plates. For the initial screening, sitting drop plates were set up with Molecular Dimensions screens JCSG Plus, PACT Premier, and MemGold at 4 °C and 21 °C. For further optimization, the best hits were set up in 24-well plates with a precipitant gradient screen. The best diffracting crystals grew in 100 mM Na Acetate pH 4.5, PEG400 15% - 22.5% at 21 °C. Diffraction data were collected at the P11 beamline at PETRAIII/DESY (Hamburg, Germany) using the standard beamline settings. The data were processed using an automated XDS processing pipeline. Initial phases were calculated in Phaser^35^ (CCP4 programme suite^36^) using PDBID:5LTR for mNG and an AlphaFold3^37^ prediction for sdAB(mNG) as search models. The crystals were of space group P2_1_2_1_2_1_ and contained single copies of mNG and sdAb(mNG) in the asymmetric unit. After initial phasing, the obtained model was iteratively refined in Phenix^38^ and manually adjusted in coot^39^. Data processing and refinement statistics are given in Supp. Table 1.

### SAXS data collection and processing

The sample used for crystallization, at a concentration of 15.2 mg/ml, was further analyzed on the P12 EMBL BioSAXS beamline (DESY, Hamburg, Germany)^40^ in SEC-SAXS mode on a Superdex 200 Increase column equilibrated in buffer containing 25 mM HEPES pH 7.6, 300 mM NaCl, 5% glycerol, 1 mM TCEP. Data were processed and analyzed using the ATSAS software suite^41^. Sample and buffer frame selection were performed in CHROMIXS^42^, the scattering curves for crystal structures were calculated in CRYSOL, and distance distributions in GNOM. *Ab initio* models were created using DAMMIN, GASBOR^43^, and DENSS^44^.

### *In vivo* Zebrafish Method

All zebrafish methods were approved by the Stony Brook University Institutional Animal Care and Use Committee. The zebrafish *tbxta-mNG* fusion line was generated using a previously published strategy^45^. CRISPR/Cas9 was used to generate a double-stranded break adjacent to the *tbxta* stop codon using a guide RNA that targets the reverse strand of the genomic sequence 5’ (CCT)GTGGCTCAGAGCTACTGAGA 3’ with the PAM site in parentheses. A repair template was generated by PCR amplification of the mNG coding sequence using primers that add 40 base pairs of homology to each side of the genomic integration site, which lies between the last amino acid codon and the stop codon of *tbxta*. The primers contain a 5’ biotin and phosphorothioate linkages in the first five bases. The PCR repair template was co-injected with Cas9 protein, guide RNA, homology-directed repair (HDR) enhancer (IDT), and 1% DMSO into 1-cell-stage zebrafish embryos. Injected embryos were sorted for mNG fluorescence and grown to adulthood to screen for germ-line transmission. For depletion experiments, an expression plasmid was designed with the *sdAb(mNG)* coding sequence fused to the zebrafish *fbxw11b* F-box protein sequence, followed by the viral p2a sequence and the mCherry coding sequence. The expression of the construct is controlled by the *hsp70l* heat-shock-inducible promoter. 25 picograms of plasmid were injected into the one-cell stage of embryos from a *tbxta-mNG* heterozygous in-cross. Control experiments were performed with a heat-shock-inducible mCherry plasmid that lacks the *fbxw11b-sdAb(mNG)* sequence. Embryos were heat-shock-induced at 6 hours post-fertilization (hpf) and imaged at 24 hpf.

## References

1. Shaner, N. C. et al. A bright monomeric green fluorescent protein derived from Branchiostoma lanceolatum. Nat Methods 10, 407–409 (2013).

2. Campbell, R. E. et al. A monomeric red fluorescent protein. Proc. Natl. Acad. Sci. 99, 7877–7882 (2002).

3. Zacharias, D. A., Violin, J. D., Newton, A. C. & Tsien, R. Y. Partitioning of lipid-modified monomeric GFPs into membrane microdomains of live cells. *Science (New York*, NY) 296, 913–916 (2002).

4. Vicidomini, R. et al. Versatile nanobody-based approach to image, track and reconstitute functional Neurexin-1 in vivo. Nat. Commun. 15, 6068 (2024).

5. Muyldermans, S. Nanobodies: Natural Single-Domain Antibodies. Annual review of biochemistry (2013) doi:10.1146/annurev-biochem-063011-092449.

6. Götzke, H. et al. The ALFA-tag is a highly versatile tool for nanobody-based bioscience applications. Nat Commun 10, 4403 (2019).

7. Sograte-Idrissi, S. et al. Circumvention of common labelling artefacts using secondary nanobodies. Nanoscale 12, 10226–10239 (2020).

8. Mougios, N. et al. NanoPlex: a universal strategy for fluorescence microscopy multiplexing using nanobodies with erasable signals. Nat. Commun. 15, 8771 (2024).

9. Unterauer, E. M. et al. Spatial proteomics in neurons at single-protein resolution. Cell 187, 1785–1800.e16 (2024).

10. Goel, R. et al. A versatile nanobody platform for live and super-resolution imaging of synaptic vesicle dynamics and plasticity in rodent and human neurons. J. Nanobiotechnology (2026) doi:10.1186/s12951-026-04489-w.

11. Rothbauer, U. et al. Targeting and tracing antigens in live cells with fluorescent nanobodies. Nature methods 3, 887–889 (2006).

12. Real, K. Q. Z. V. et al. A Versatile Synaptotagmin-1 Nanobody Provides Perturbation-Free Live Synaptic Imaging And Low Linkage-Error in Super-Resolution Microscopy. Small Methods 7, e2300218 (2023).

13. Shen, F. & Dassama, L. M. K. Opportunities and challenges of protein-based targeted protein degradation. Chem. Sci. 14, 8433–8447 (2023).

14. Daniel, K. et al. Conditional control of fluorescent protein degradation by an auxin-dependent nanobody. Nat. Commun. 9, 3297 (2018).

15. Borg, S. et al. An Intracellular Nanotrap Redirects Proteins and Organelles in Live Bacteria. mBio 6, 10.1128/mbio.02117-14 (2015).

16. Gerdes, C. et al. A nanobody-based fluorescent reporter reveals human α-synuclein in the cell cytosol. Nat. Commun. 11, 2729 (2020).

17. Cheloha, R. W. et al. Improved GPCR ligands from nanobody tethering. Nat. Commun. 11, 2087 (2020).

18. Feng, S. et al. Improved split fluorescent proteins for endogenous protein labeling. Nat. Commun. 8, 370 (2017).

19. Robust fluorescent proteins for high-resolution microscopy and biochemical techniques. Nat Methods 1–2 (2022) doi:10.1038/s41592-022-01661-6.

20. Bindels, D. S. et al. mScarlet: a bright monomeric red fluorescent protein for cellular imaging. Nat. Methods 14, 53–56 (2017).

21. Mohr, D., Frey, S., Fischer, T., Güttler, T. & Görlich, D. Characterisation of the passive permeability barrier of nuclear pore complexes. EMBO J. 28, 2541–2553 (2009).

22. Schulte-Merker, S., Eeden, F. J. M. van, Halpern, M. E., Kimmel, C. B. & Nüsslein-Volhard, C. no tail (ntl) is the zebrafish homologue of the mouse T (Brachyury) gene. Development 120, 1009–1015 (1994).

23. Halpern, M. E., Ho, R. K., Walker, C. & Kimmel, C. B. Induction of muscle pioneers and floor plate is distinguished by the zebrafish no tail mutation. Cell 75, 99–111 (1993).

24. Yamaguchi, N., Colak-Champollion, T. & Knaut, H. zGrad is a nanobody-based degron system that inactivates proteins in zebrafish. eLife 8, e43125 (2019).

25. Shamsudin, Y., Walker, A. R., Jones, C. M., Martínez, T. J. & Boxer, S. G. Simulation-guided engineering of split GFPs with efficient β-strand photodissociation. Nat. Commun. 14, 7401 (2023).

26. Köker, T., Fernandez, A. & Pinaud, F. Characterization of Split Fluorescent Protein Variants and Quantitative Analyses of Their Self-Assembly Process. Sci. Rep. 8, 5344 (2018).

27. Ries, J., Kaplan, C., Platonova, E., Eghlidi, H. & Ewers, H. A simple, versatile method for GFP-based super-resolution microscopy via nanobodies. Nat. Methods 9, 582–584 (2012).

28. D’Este, E., Lukinavičius, G., Lincoln, R., Opazo, F. & Fornasiero, E. F. Advancing cell biology with nanoscale fluorescence imaging: essential practical considerations. Trends Cell Biol. (2024) doi:10.1016/j.tcb.2023.12.001.

29. Bloch, J. S. et al. Development of a universal nanobody-binding Fab module for fiducial-assisted cryo-EM studies of membrane proteins. Proc. Natl. Acad. Sci. 118, e2115435118 (2021).

30. Laverty, D. et al. Cryo-EM structure of the human α1β3γ2 GABAA receptor in a lipid bilayer. Nature 565, 516–520 (2019).

31. Shaib, A. H. et al. One-step nanoscale expansion microscopy reveals individual protein shapes. Nat. Biotechnol. 1–9 (2024) doi:10.1038/s41587-024-02431-9.

32. Bradbury, A. & Plückthun, A. Reproducibility: Standardize antibodies used in research. Nature 518, 27–29 (2015).

33. Maidorn, M., Olichon, A., Rizzoli, S. O. & Opazo, F. Nanobodies reveal an extra-synaptic population of SNAP-25 and Syntaxin 1A in hippocampal neurons. mAbs 11, 305–321 (2018).

34. Keihani, S. et al. The long noncoding RNA neuroLNC regulates presynaptic activity by interacting with the neurodegeneration-associated protein TDP-43. Sci Adv 5, eaay2670 (2019).

35. McCoy, A. J. et al. Phaser crystallographic software. J Appl Crystallogr 40, 658–674 (2007).

36. Martinez-Ripoll, M. & Albert, A. Continuous development in macromolecular crystallography with CCP4. Acta Crystallogr. Sect. D 79, 447–448 (2023).

37. Abramson, J. et al. Accurate structure prediction of biomolecular interactions with AlphaFold 3. Nature 630, 493–500 (2024).

38. Liebschner, D. et al. Macromolecular structure determination using X-rays, neutrons and electrons: recent developments in Phenix. Acta Crystallogr Sect D 75, 861–877 (2019).

39. Emsley, P., Lohkamp, B., Scott, W. G. & Cowtan, K. Features and development of Coot. Acta Crystallogr Sect D Biological Crystallogr 66, 486–501 (2010).

40. Blanchet, C. E. et al. Versatile sample environments and automation for biological solution X-ray scattering experiments at the P12 beamline (PETRA III, DESY). J. Appl. Crystallogr. 48, 431–443 (2015).

41. Manalastas-Cantos, K. et al. ATSAS 3.0: expanded functionality and new tools for small-angle scattering data analysis. J. Appl. Crystallogr. 54, 343–355 (2021).

42. Panjkovich, A. & Svergun, D. I. CHROMIXS: automatic and interactive analysis of chromatography-coupled small-angle X-ray scattering data. Bioinformatics 34, 1944–1946 (2017).

43. Svergun, D. I., Petoukhov, M. V. & Koch, M. H. J. Determination of Domain Structure of Proteins from X-Ray Solution Scattering. Biophys. J. 80, 2946–2953 (2001).

44. Grant, T. D. Ab initio electron density determination directly from solution scattering data. Nat. Methods 15, 191–193 (2018).

45. Zhang, Y., Marshall-Phelps, K. & Almeida, R. G. Fast, precise and cloning-free knock-in of reporter sequences in vivo with high efficiency. Development 150, dev201323 (2023).

