## Supplementary Figure for "A High-Affinity Nanobody Recognizing mNeonGreen Enables Versatile Biochemical, Cellular, and *in vivo* Applications"

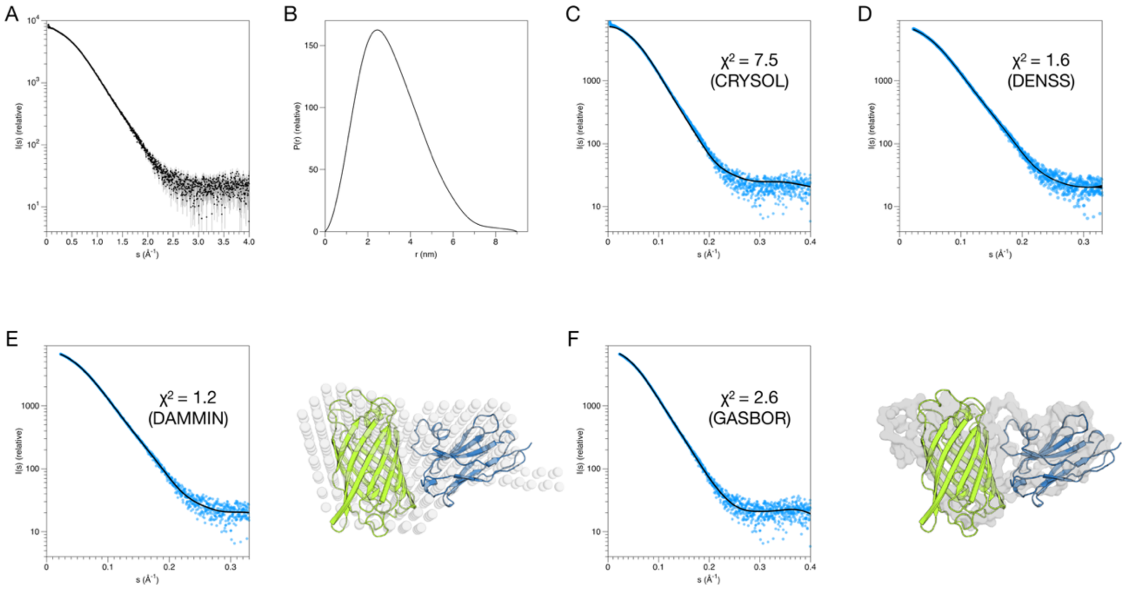

**Supplementary Figure 1. Validation of the crystal structure in solution using SAXS.**
**A.** Raw scattering data for the complex. **B.** The distance distribution function shows a maximum distance of 9 nm. **C.** Fit of the crystal structure (black line) to the SAXS data (blue dots). **D.** Electron density reconstitution in DENSS. Overlaid fit of 20 independent runs (black lines) to the SAXS data. See Supplementary Movie 1 for the reconstituted electron density in solution. **E.** *Ab initio* bead-based modeling of the complex in DAMMIN. The model is shown as spheres on top of the crystal structure (cartoons). **F.** *Ab initio* chain-like modeling in GASBOR. The SAXS model is shown as a gray surface.

```
mNeonGreen      MVSKGEEDNMASLPAT--HELHIFGSINGVDFDMVGQGTGNPNDGYEELNLKSTKG-DLQ 57
mTagBFP         ---MSE----ELIKENMHMKLYMEGTVDNHHFKCTSEGEGKPYEGTQTMRIKVVEGGPLP 53
mEGFP           MVSKGEELFTGVVP----ILVELDGDVNGHKFSVSGEGEGDATYGKLTLKFICTTG-KLP 55
mCherry         MVSKGEEDNMAIIKEFMRFKVHMEGSVNGHEFEIEGEGEGRPYEGTQTAKLKVTKGGPLP 60
mRuby3          MVSKGE----ELIKENMRMKVVMEGSVNGHQFKCTGEGEGRPYEGVQTMRIKVIEGGPLP 56
Dendra          ---------MNLIKEDMRVKVHMEGNVNGHAFVIEGEGKGKPYEGTQTANLTVKEGAPLP 51
Dronpa          ---------MSVIKPDMKIKLRMEGAVNGHPFAIEGVGLGKPFEGKQSMDLKVKEGGPLP 51
mEos3.2         ---------MSAIKPDMKIKLRMEGNVNGHHFVIDGDGTGKPFEGKQSMDLEVKEGGPLP 51
mBaoJin         MVSKGEEENMASTP----FKFQIKGTINGKSFTVEGEGEGNSHEGSHKGKYVCTSG-KLP 55

mNeonGreen      FSPWILVPHIGYGFHQYLPYPDGMS--PFQAAMVDGSGYQVHRTMQFEDGASLTVNYRYT 115
mTagBFP         FAFDILATSFLYGSKTFINHTQGIP--DFFKQS-FPEGFTWERVTTYEDGGVLTATQDTS 110
mEGFP           VPWPTLVTTLTYGVQCFSRYPDHMKQHDFFKSA-MPEGYVQERTIFFKDDGNYKTRAEVK 114
mCherry         FAWDILSPQFMYGSKAYVKHPADIP--DYLKLS-FPEGFKWERVMNFEDGGVVTVTQDSS 117
mRuby3          FAFDILATSFMYGSRTFIKYPADIP--DFFKQS-FPEGFTWERVTRYEDGGVVTVTQDTS 113
Dendra          FSYDILTTAVHYGNRVFTKYPEDIP--DYFKQS-FPEGYSWERTMTFEDKGICTIRSDIS 108
Dronpa          FAYDILTTVFCYGNRVFAKYPENIV--DYFKQS-FPEGYSWERSMNYEDGGICNATNDIT 108
mEos3.2         FAFDILTTAFHYGNRVFAKYPDNIQ--DYFKQS-FPKGYSWERSLTFEDGGICNARNDIT 108
mBaoJin         MSWAALGTTFGYGMKYYTKYPSGLK--NWFREV-MPGGFTYDRHIQYKGDGSIHAKHQHF 112

mNeonGreen      YEGSHIKGEAQVKGTGFPADGPVMTNSLTAADWCR-SKKTYPN---DKTIISTFKWSYT- 170
mTagBFP         LQDGCLIYNVKIRGVNFTSNGPVMQKKT--LGWEAFTETLYPA---DGGLEGRNDMALK- 164
mEGFP           FEGDTLVNRIELKGIDFKEDGNILGHKLE--YNYNSHNVYIMADKQKNGIKVNFKIRHN- 171
mCherry         LQDGEFIYKVKLRGTNFPSDGPVMQKKT--MGWEASSERMYPE---DGALKGEIKQRLK- 171
mRuby3          LEDGELVYNVKVRGVNFPSNGPVMQKKT--KGWEPNTEMMYPA---DGGLRGYTDIALK- 167
Dendra          LEGDCFFQNVRFKGTNFPPNGPVMQKKT--LKWEPSTEKLHVR---DGLLVGNINMALL- 162
Dronpa          LDGDCYIYEIRFDGVNFPANGPVMQKRT--VKWEPSTEKLYVR---DGVLKGDVNMALS- 162
mEos3.2         MEGDTFYNKVRFYGTNFPANGPVMQKKT--LKWEPSTEKMYVR---DGVLTGDIEMALL- 162
mBaoJin         MKNGTYHNIVEFTGQDFKENSPVLTGDMN--VSLPNEVPQIPR---DDGVECPVTLLYPL 167

mNeonGreen      -TGNGKRYRSTARTTYTFAKPMAANYLKNQ--PMYVFRKTELKHSKTELNFKEWQKAFT- 226
mTagBFP         -LVGGSHLIANIKTTYRSKKPAKNLKMPGV--YYVDYRLERIKEANNETYVEQHEVAVAR 221
mEGFP           -IEDGSVQL---ADHYQQNTPIGDGPVLLPDNHYLSTQSK-LSKDPNEKRDHMVLLEFVT 226
mCherry         -LKDGGHYDAEVKTTYKAKKPV---QLPGA--YNVNIKLDITSHNEDYTIVEQYERAE-- 223
mRuby3          -VDGGGHLHCNFVTTYRSKKTVGNIKMPGV--HAVDHRLERIEESDNETYVVQREVAVAK 224
Dendra          -LEGGGHYLCDFKTTYKAK-KV--VQLPDA--HFVDHRIEILGNDSDYNKVKLYEHAVAR 216
Dronpa          -LEGGGHYRCDFKTTYKAK-KV--VQLPDY--HFVDHHIEIKSHDKDYSNVNLHEHAEAH 216
mEos3.2         -LEGNAHYRCDFRTTYKAKEKG--VKLPGA--HFVDHCIEILSHDKDYNKVKLYEHAVAH 217
mBaoJin         LSDKSKYVE---AHQYTICKPLHNQPAPDVPYHWIRKQYT-QSKDDAEERDHICQSETL- 222

mNeonGreen      ---DVMGMDELYK 236
mTagBFP         YCDLPSKLGHKLN 234
mEGFP           AAGITLGMDELYK 239
mCherry         GRHSTGGMDELYK 236
mRuby3          YSNLGGGMDELYK 237
Dendra          YSPLPSQAW---- 225
Dronpa          S-ELPRQAK---- 224
mEos3.2         S-GLPDNARR--- 226
mBaoJin         -EAHLKGMDELYK 234
```

**Supplementary Figure 2. Sequence alignment of fluorescent proteins used in IP experiments.**

Positions at which mNeonGreen interacts directly with sdAb(mNG) are indicated with blue boxes.

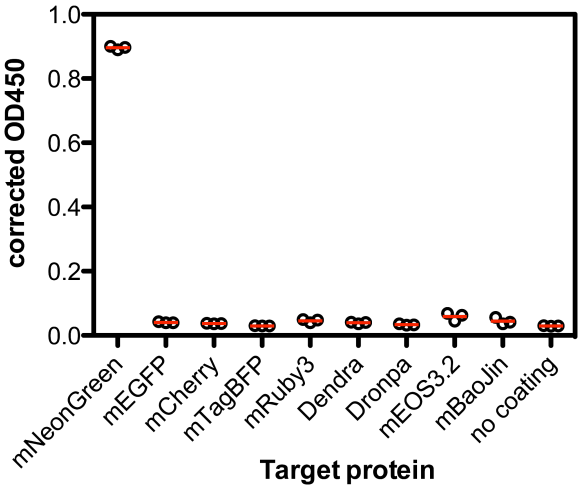

**Supplementary Figure 3. Specificity of sdAb(mNG) for diverse classes of fluorescent proteins**.
For the ELISA, 100 fmol of target protein was coated in a single well and detected with 3 nM biotinylated sdAb (mNG), followed by Streptavidin-HRP. Raw absorbance values at 450 nm are given as black circles, average values are denoted as red lines.

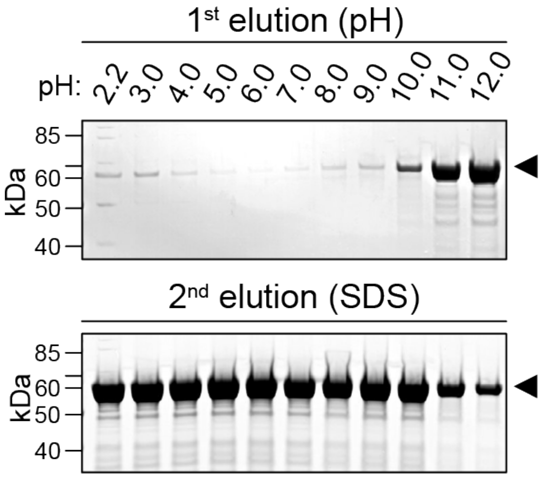

**Supplementary Figure 4. pH-dependent elution of the mNeonGreen Selector**.

To assess the elution of the mNeonGreen Selector at different pH, the resin was saturated with recombinantly expressed and affinity-purified mNeonGreen. Excess protein was removed by extensive washing, after which the resin was incubated in buffer solutions at the indicated pH values, followed by a second elution using hot SDS sample buffer. Eluates were analyzed by SDS-PAGE. **Top**: First elution of the Selector resin at the indicated pH. **Bottom**: Second elution of the Selector resin using SDS sample buffer. Each lane represents an individual experiment performed under the indicated conditions. Arrowheads point to the target protein.

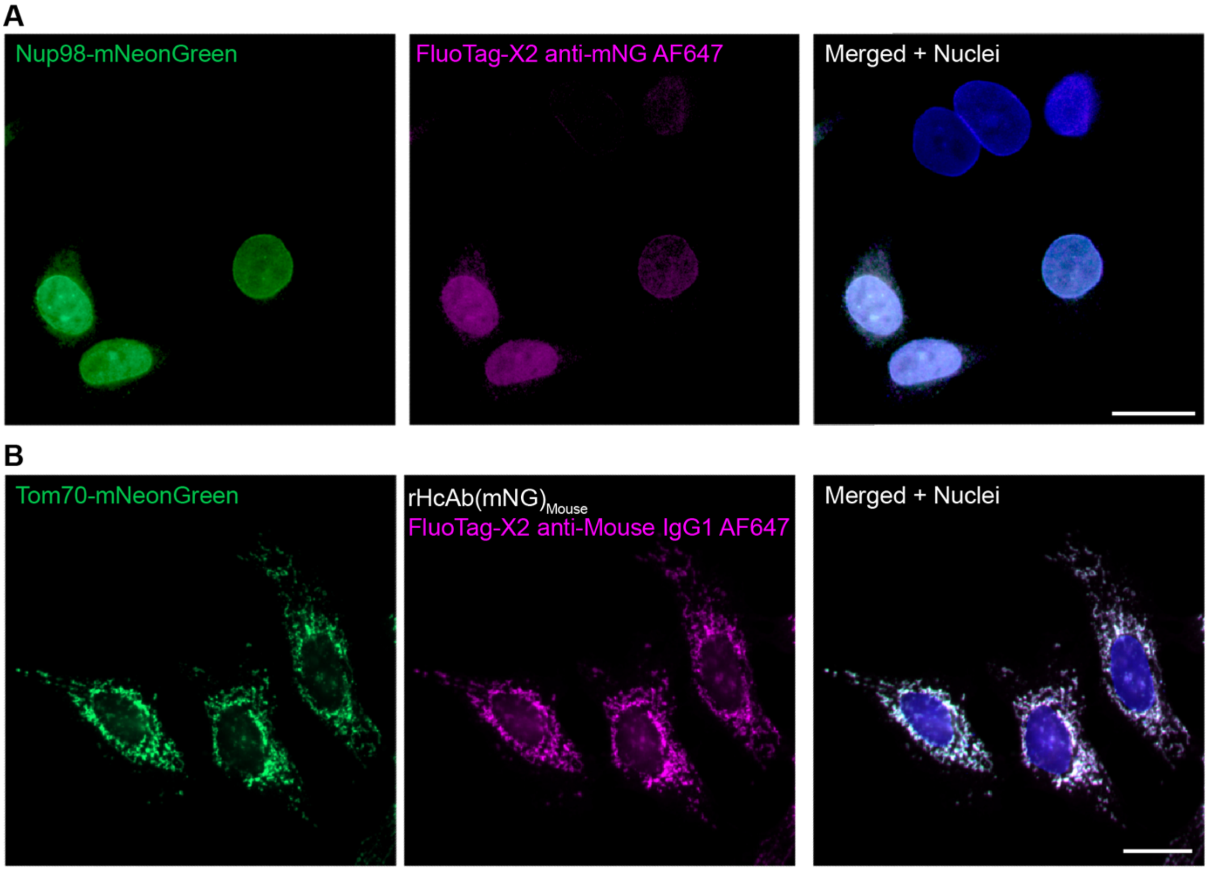

**Supplementary Figure 5: Detection of mNeonGreen-fusion proteins using sdAb(mNG) in mono- and bivalent formats.**

**A:** HeLa cells were transiently transfected with Nup98-mNG (green) and fixed with 4% PFA before detecting mNG with the monovalent reagent FluoTag®-X2 anti-mNeonGreen AF647 (magenta). **B**: HeLa cells were transiently transfected with TOM70-mNG (green) and fixed with 4% PFA. mNG was detected with the bivalent reagent sdAb(mNG) fused to a mouse IgG Fc domain (recombinant heavy chain only antibody; $rHcAb(mNG)_{Mouse}$) as primary antibody and FluoTag®-X2 anti-mouse-IgG1 as secondary antibody. **Left**: mNeonGreen fluorescence signal (green). **Middle**: Fluorescence signal obtained using the respective anti-mNG reagent in direct or indirect staining (magenta). **Right**: Merge of the images displayed, including an anti-Lamin staining (blue) to visualize nuclei. Scale bar: 20 µm.

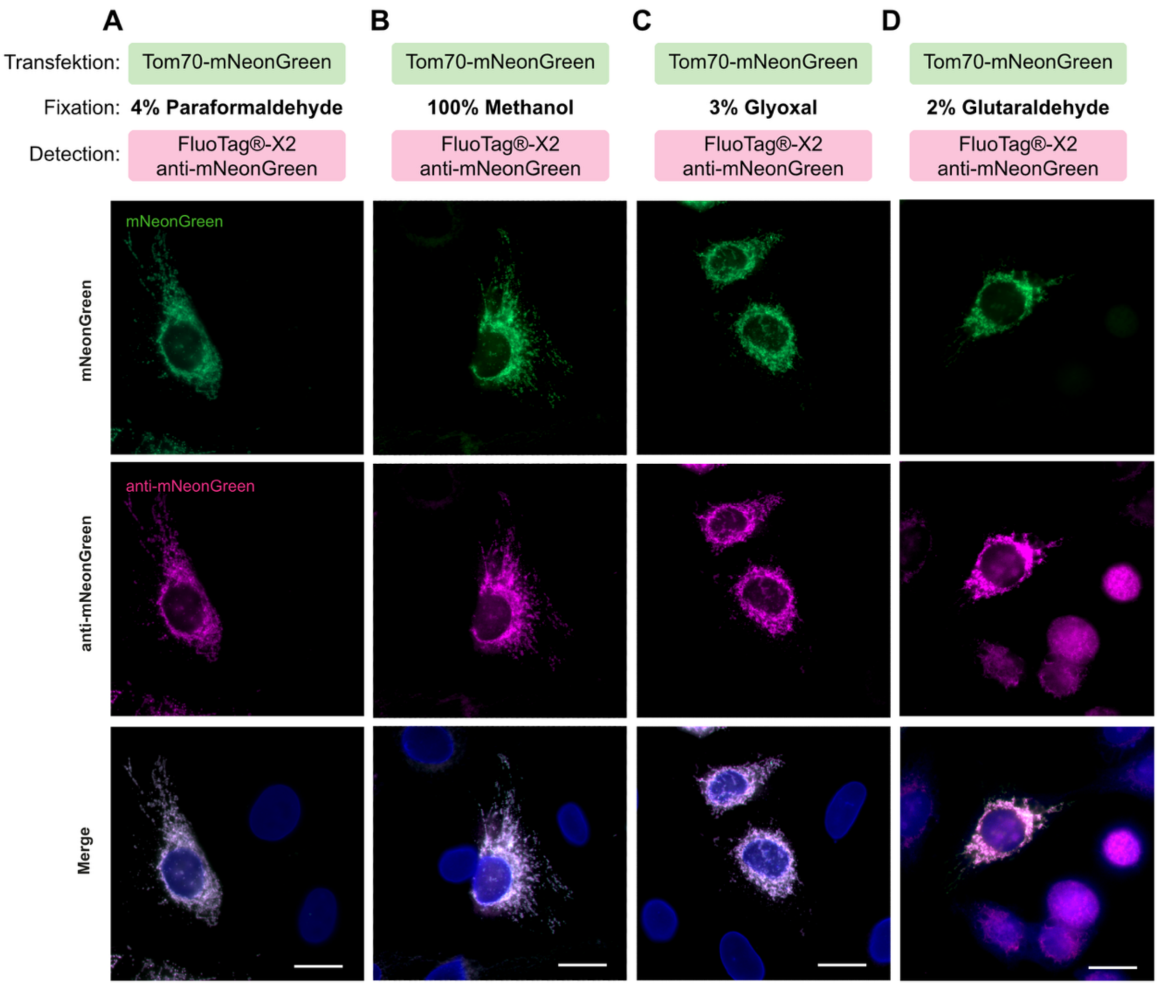

**Supplementary Figure 6: mNeonGreen recognition by sdAb(mNG) is preserved across fixation methods.**

HeLa cells were transiently transfected with Tom70-mNG and fixed using 4% Paraformaldehyde (**A**), 100% Methanol (**B**), 3% Glyoxal (**C**) or 2% Glutaraldehyde (**D**), followed by staining with FluoTag®-X2 anti-mNG AF568. All samples were counterstained for LaminA/C. Shown are mNG fluorescence (top, green), anti-mNG signal (middle, magenta) and merged images including anti-LaminA/C counterstaining in blue (bottom). Scale bar: 20 µm.

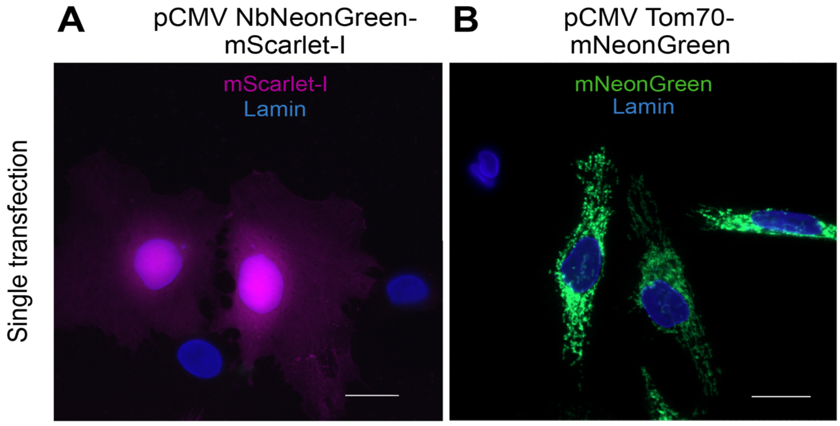

**Supplementary Figure 7**. Related to Fig. 7

Controls with single plasmid transfections imaged under an epifluorescence microscope. Signal distribution on HeLa cells transiently transfected with plasmids encoding sdAb(mNG)-mScarlet-I (**A**) or Tom70-mNG (**B**). Scale bar: 20µm

**Supplementary Table 1.** X-ray data processing and refinement statistics. The values in parentheses correspond to the highest-resolution shell.

| | **mNG / sdAB(mNG) PDBID: 30TF** |
|---|---|
| **Data processing** | |
| Space group | $P\ 2_1\ 2_1\ 2_1$ |
| Unit cell | |
| a b c (Å) | 53.662 65.308 135.041 |
| α β γ (°) | 90 90 90 |
| Resolution range (Å) | 50 – 1.26 (1-34 – 1.26) |
| ⟨I/σI⟩ | 21.7 (1.1) |
| Completeness (%) | 96.2 (80.5) |
| Redundancy | 12.6 (8.7) |
| $R_{meas}$ (%) | 5.8 (169.7) |
| $CC_{1/2}$ (%) | 100 (58.5) |
| **Structure refinement** | |
| $R_{work}/R_{free}$ (%) | 12.3 / 14.4 |
| RMSD bond lengths (Å) | 0.020 |
| RMSD bond angles (°) | 1.6 |
| Rama-Z score | |
| whole | 1.00 (0.40) |
| helix | -1.32 (1.03) |
| sheet | 0.82 (0.36) |
| loop | 1.02 (0.43) |
| Molprobity score | 1.54 |
| Clashscore | 7.33 |
| Rotamer outliers (%) | 1.5 |
| Ramachandran (%) | |
| favoured | 98.81 |
| allowed | 1.19 |
| outliers | 0 |

**Supplementary Table 2:** Plasmids for eukaryotic expression used in this study

| **Plasmid Name** | **Encoded construct** | **Used in Figure** | **Source** |
|---|---|---|---|
| pNT1705 | TOM70-mNG | 5 & Supp. 5, 6, 7 | This paper |
| pNT4316 | sdAb(mNG)-mScarlet-I | 7 | This paper |
| pNT4596 | Tom70-mNG-ALFA | 7 | This paper |
| pNT4594 | mNG | 7 | This paper |
| pNT4727 | NES-sdAb(mNG)-ALFA | 7 | This paper |
| pNT4595 | NLS-mNG | 7 | This paper |
| pNT4599 | Nup98-mNG | Supp. 5 | This paper |
| iNbmNG-mScarlet_AAV | sdAb(mNG)-mScarlet | 8 & Supp. Movie2/3 | This paper |
| pmNeonGreenHO-G | Soluble mNG | 6 | gift from Isei Tanida Addgene # 127912 |
| HsTPC2-mNeonGreen | human TPC2 tagged with mNG at C-terminus | 6 | This paper |
| $NG2_{11}$-miRFP670nano3 | Cytosolic miRFP670nano3 tagged at N-terminus with the single beta-strand 11 of mNG2 | 6 | This paper |
| HsTPC2-$mNG2_{1-10}$-miRFP670nano3 | human TPC2 tagged with a $mNG2_{(1-10)}$ and miRFP670nano3 at its cytosolic C-terminus | 6 | This paper |

**Supplementary Table 3:** Antibodies used in this study.

| Target | Description/Name | Used in Figure | dilution / conc. | Company | Cat. Number | RRID |
|---|---|---|---|---|---|---|
| mNeonGreen | sdAb anti-mNeonGreen (1E2) | 1, 2 | various | NanoTag Biotechnologies | N3205 | AB_3076073 |
| mNeonGreen | Rabbit IgG Fc-fusion sdAb (mNG) (rHcAb(mNG)$_{Rabbit}$) | 5B | 1:500 / 0.2 mg/ml | NanoTag Biotechnologies | N3283 | Pending |
| mNeonGreen | FluoTag®-X2 anti-mNeonGreen Alexa Fluor™ 647 | 5A | 1:500 / 10 nM | NanoTag Biotechnologies | N3202-AF647-L | AB_3076070 |
| mNeonGreen | FluoTag®-X2 anti-mNeonGreen Atto 643 | 5C | 1:500 / 10 nM | NanoTag Biotechnologies | N3202-At643-L | AB_3076072 |
| mNeonGreen | FluoTag®-X2 anti-mNeonGreen Alexa Fluor™ 568 | 6, Supp. 6 | 1:500 / 10 nM | NanoTag Biotechnologies | N3202-AF568-L | Pending |
| Rabbit IgG | FluoTag®-X2 anti-Rabbit IgG Alexa Fluor™ 647 | 5B | 1:500 / 20 nM | NanoTag Biotechnologies | N2402-AF647-S | AB_3076036 |
| ALFA-tag | FluoTag®-X2 anti-ALFA® Alexa Fluor™ 647 | 7B | 1:500 / 10 nM | NanoTag Biotechnologies | N1502-AF647-L | AB_3075981 |
| Mouse IgG1 | FluoTag®-X2 anti-Mouse IgG1 Alexa Fluor™ 647 | Supp. 5 | 1:500 / 20 nM | NanoTag Biotechnologies | N2002-AF647-S | AB_3076020 |
| mNeonGreen | Mouse IgG1 Fc-fusion sdAb (mNG) (rHcAb(mNG)$_{Mouse}$) | Supp. 5 | 1:500 / 0.2 mg/ml | NanoTag Biotechnologies | N3282 | Pending |
| Mouse IgG2 | FluoTag®-X2 anti-Mouse IgG2 Alexa Fluor™ 568 | 5, 6, Supp. 5 | 1:500 / 0.2 mg/ml | NanoTag Biotechnologies | N2702-AF568-S | Pending |
| Mouse IgG2 | FluoTag®-X2 anti-Mouse IgG2 Alexa Fluor™ 647 | Supp. 6, 7 | 1:500 / 0.2 mg/ml | NanoTag Biotechnologies | N2702-AF568-S | AB_2936181 |
| LaminA/C | anti-LaminA/C (Mouse IgG2) | 5, 6, Supp. 5-7 | 1:2000 / 0.1 mg/ml | AbCam | ab238303 | AB_3722722 |

**Supplementary Movie 1.** SAXS-based electron density reconstitution of the complex in solution. The crystal structure is overlaid as a cartoon.

**Supplementary Movie 2.** Live hippocampal rat neurons imaged under epifluorescence microscopy that were co-infected with NbSyt1-mNG (green signal) and sdAb(mNG)-mScarlet-I (magenta signal).

**Supplementary Movie 3.** Live hippocampal rat neurons imaged under epifluorescence microscopy that were co-infected with NbSyt1-mNG (green signal) and sdAb(mNG)-mScarlet-I (magenta signal).
